# Non-productive exposure of PBMCs to SARS-CoV-2 induces cell-intrinsic innate immunity responses

**DOI:** 10.1101/2022.02.15.480527

**Authors:** Julia Kazmierski, Kirstin Friedmann, Dylan Postmus, Cornelius Fischer, Jenny Jansen, Anja Richter, Laure Bosquillon de Jarcy, Christiane Schüler, Madlen Sohn, Sascha Sauer, Christian Drosten, Antoine-Emmanuel Saliba, Leif Erik Sander, Daniela Niemeyer, Christine Goffinet

## Abstract

Cell-intrinsic responses mounted *in vivo* in PBMCs during mild and severe COVID-19 differ quantitatively and qualitatively. Whether they are triggered by signals emitted by productively infected cells of the respiratory tract or are, at least partially, resulting from physical interaction with virus particles, remains unclear. Here, we analyzed susceptibility and expression profiles of PBMCs from healthy donors upon *ex vivo* exposure to SARS-CoV and SARS-CoV-2. In line with the absence of detectable ACE2 receptor expression, human PBMCs were refractory to productive infection. Bulk and single cell RNA-sequencing revealed JAK/STAT-dependent induction of interferon-stimulated genes, but not pro-inflammatory cytokines. This SARS-CoV-2-specific response was most pronounced in monocytes. SARS-CoV-2-RNA-positive monocytes displayed a lower ISG signature as compared to bystander cells of the identical culture. This suggests a preferential invasion of cells with a low ISG base-line profile or delivery of a SARS-CoV-2-specific sensing antagonist upon efficient particle internalization. Together, non-productive physical interaction of PBMCs with SARS-CoV-2-but not SARS-CoV particles stimulates JAK/STAT-dependent, monocyte-accentuated innate immune responses that resemble those detected *in vivo* in patients with mild COVID-19.

## Introduction

The current SARS-CoV-2 pandemic represents a global medical, societal and economical emergency of increasing importance. Arising at the end of 2019 in the Hubei province in China, the causative agent of the coronavirus disease 2019 (COVID-19), SARS-CoV-2, has to date infected more than 395 million individuals world-wide (World Health Organization). Owing to SARS-CoV-2 infection, more than 5,7 million deaths were reported up to today (as of 2022, February 6th). The predominant symptoms of symptomatic COVID-19 are fever, cough, and shortness of breath, however, in severe cases disease can progress to pneumonia, acute respiratory distress syndrome, and multiple organ failure (Chen et al. 2020; Wölfel et al. 2020). The management of the pandemic is complicated by a relatively low manifestation index, a large inter-individual spectrum of clinical courses ranging from asymptomatic to fatal outcomes, pre- and asymptomatic infectious phases (Jones et al. 2021; Rothe et al. 2020), and the ongoing emergence of variants with increased transmissibility and/or immune escape. The reasons for the high inter-individual outcome of infection are insufficiently understood and may include different degrees of cross-reactive background immunity at the level of humoral (Anderson et al. 2021; K. W. Ng et al. 2020) and T-cell-mediated immunity (Braun et al. 2020; Bacher et al. 2020; Nelde et al. 2021; Schulien et al. 2021), polymorphisms in genes related to innate immunity (Zhang et al. 2020) and autoimmunity (Bastard et al. 2020). Currently, specific treatment regimen must be administered early post-infection. They include the RNA polymerase inhibitor Remdesivir that may reduce hospitalization time but not mortality (Y. Wang et al. 2020) and monoclonal anti-spike antibodies with variant-specific neutralization potencies (Weinreich et al. 2021; RECOVERY Collaborative Group, Horby, Mafham, et al. 2021). In the late phase of infection, the administration of the immune modulator dexamethasone (RECOVERY Collaborative Group, Horby, Lim, et al. 2021) dampens hyperactivation of cytokine-driven immune responses. While several effective vaccines are available, the necessity for specific treatment options will likely persist given the expected proportion of the population that will not have access to vaccines or will refuse vaccination.

To accelerate the establishment of immunomodulatory strategies, it is crucial to characterize *ex vivo* systems that correlate with cellular immunophenotypes of SARS-CoV-2 infection *in vivo* and that may contribute to pre-clinical testing. Furthermore, the usage of *ex vivo* platforms allows the systematic and comparative investigation of human cellular responses to exposure with different representatives of the species SARS-related coronaviruses, including SARS-CoV. Peripheral immune cells are major contributors to human cellular responses upon infection. Given the recruitment of blood mononuclear cells to the lung compartment (Delorey et al. 2021; Bost et al. 2020; Wendisch et al. 2021), and the reported presence of viral RNA detectable in the peripheral blood of up to 10% severely ill patients (Prebensen et al. 2021; Andersson et al. 2020), direct contact of PBMCs with SARS-CoV-2 virions is a likely scenario.

Here, we analyzed susceptibility to infection and cell-intrinsic innate responses of peripheral blood cells from healthy donors upon *ex vivo* exposure to SARS-CoV and SARS-CoV-2. Although both SARS-related coronaviruses failed to detectably replicate and spread in PBMCs, SARS-CoV-2 specifically triggered a JAK/STAT-dependent innate immune response that was most pronounced in monocytes. Single-cell, virus-inclusive RNA sequencing revealed relatively inefficient and ACE2-independent uptake of virus particles and a SARS-CoV-2 exposure-specific gene expression profile. Cellular responses, consisting in upregulation of expression of interferon-stimulated genes (ISGs) but not pro-inflammatory cytokines, partially recapitulate expression profiles obtained by single-cell RNA-sequencing of PBMCs from patients experiencing mild COVID-19 (Arunachalam et al. 2020; Schulte-Schrepping et al. 2020; Silvin et al. 2020). Our data demonstrate that cells from the peripheral blood, when undergoing contact to SARS-CoV-2 particles, mount cellular responses that potentially contribute to control and/or pathogenesis of the infection.

## RESULTS

### Absence of productive infection of human PBMCs by SARS-CoV and SARS-CoV-2

To address the ability of SARS-related coronaviruses to infect and propagate in cells of the peripheral blood, we exposed unstimulated PBMCs from healthy individuals to purified stocks of SARS-CoV and SARS-CoV-2, respectively, using equal infectious titers as determined on Vero E6 cells. As a reference, PBMCs were exposed to supernatants from uninfected Vero E6 cells (mock-exposed). For both SARS-related coronaviruses, infectivity in cell culture supernatants drastically decreased over time compared to the inoculum, reaching undetectable levels at three days post-inoculation (**Fig. 1A**), pointing towards absence of *de novo* production of infectious particles. Treatment of cells with the polymerase inhibitor Remdesivir did not further reduce infectivity in the supernatant, suggesting that the infectivity detectable at 24 hours post-inoculation reflects virus input (**Fig. 1B**). In contrast, infection of Vero E6 cells with the identical SARS-CoV-2 stock gave rise to a productive and Remdesivir-sensitive infection (**Supplemental Fig. 1**). In our experiments, virus-containing supernatant was replaced with fresh medium four hours post-inoculation. Nevertheless, viral RNA genome equivalents remained detectable in the culture supernatant until the end of the experiment for both SARS-CoV and SARS-CoV-2 (up to 192 hours post-exposure) (**Fig. 1C**). Viral RNA was abundant also in supernatants from Remdesivir-treated cultures and cultures exposed to heat-inactivated SARS-CoV-2 until 192 hours post-exposure, arguing for a high stability of the residual viral RNA of the inoculum and against a constant replenishment of extracellular viral RNA pools as a reason for the stable RNA quantities (**Fig. 1D**), in line with reported longevity of the incoming genomic viral RNA (Lee et al. 2022). Notably, blunting signaling by type I interferons (IFNs) through constant presence of the JAK/STAT inhibitor Ruxolitinib failed to enable secretion of infectious particles and viral RNA in the supernatant, suggesting that JAK/STAT-dependent cell-intrinsic innate immunity is not the underlying reason for the absence of detectable virus production (**Fig. 1 A, C**).

**Figure 1.**
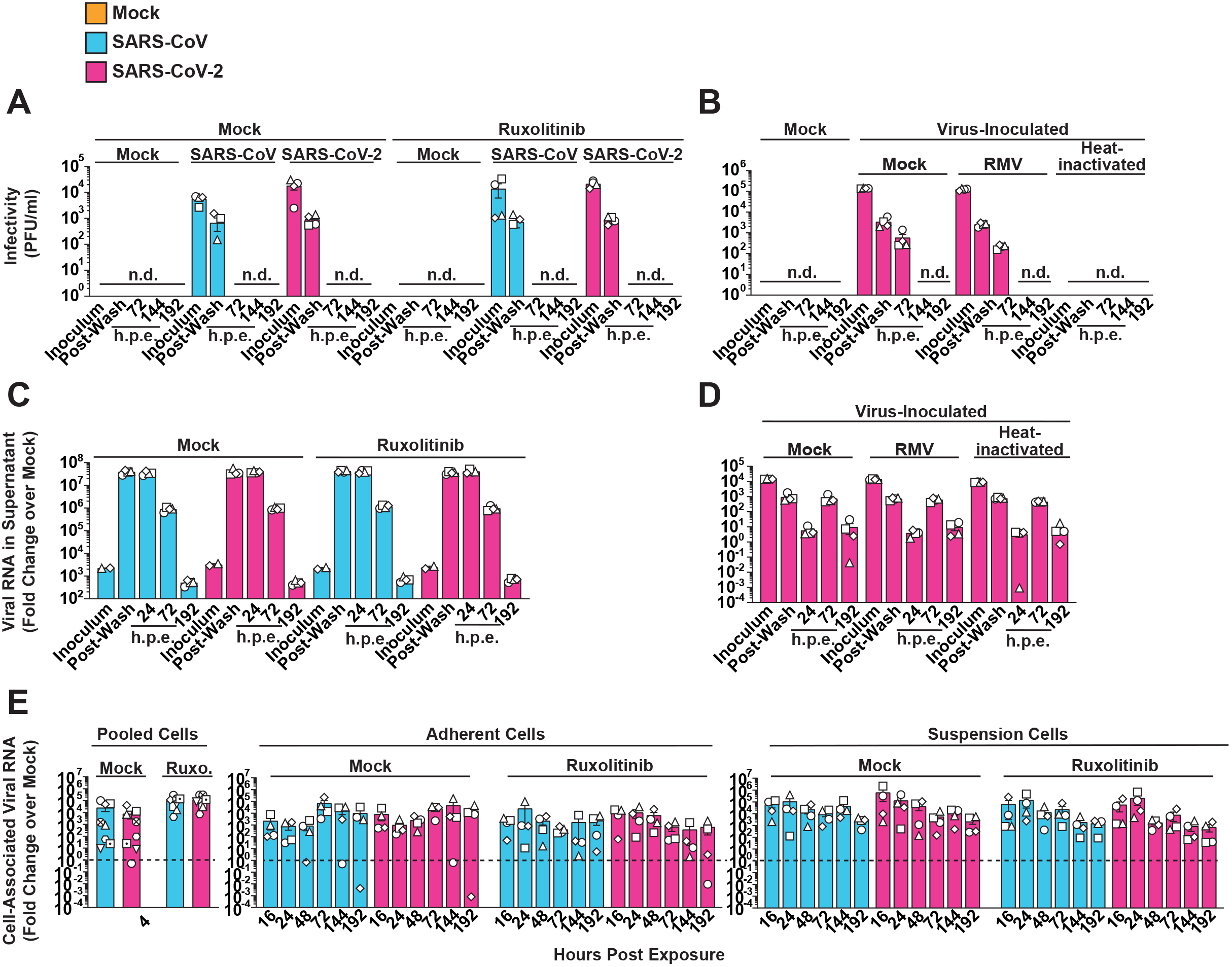
Absence of productive infection of PBMCs by SARS-CoV and SARS-CoV-2. Untreated or Ruxolitinib (10 µM)-treated PBMCs from four individual donors were exposed to SARS-CoV or SARS-CoV-2 (MOI 0.5). PBMCs inoculated with supernatant from Vero E6 cell cultures mixed with PBS and OptiPro serum-free medium supplemented with 0.5% gelatine were used as control condition (Mock). Supernatants and individual cell fractions were collected at indicated time points post-inoculation and analyzed for: (**A-B**) Infectivity in cell culture supernatants by plaque titration assay (**C-D**) Relative changes of viral RNA (genome equivalents) quantities in cell culture supernatants by Q-RT-PCR (**E**) Relative changes of cell-associated viral genomic RNA quantities by Q-RT-PCR normalized to cellular *RNASEP* expression Data were generated in four individual experiments using cells from four individual donors represented by different symbols. n.d. = not detectable; h.p.e. = hours post-exposure; RMV = Remdesivir; Ruxo. = Ruxolitinib.

To elucidate if PBMCs, despite being non-permissive, are nevertheless susceptible to SARS-related coronavirus entry and initial RNA replication, we monitored cell-associated viral RNA species in the adherent and the suspension cell fractions of the cultures over time. Cell-associated viral genome equivalents (**Fig. 1E**) and subgenomic viral E RNA (**Supplemental Fig. 2**), the latter produced during discontinuous viral transcription, remained stable over time and did not differ in quantities for both SARS-related coronaviruses. Ruxolitinib treatment did not detectably facilitate RNA replication (**Fig. 1E**, **Supplemental Fig. 2**), suggesting absence of essential cofactors at the level of entry and/or RNA replication rather than the antiviral activity of IFN-regulated restriction factors. In line with this idea, we failed to detect expression of the SARS-coronavirus receptor, angiotensin-converting enzyme 2 (ACE2) in PBMCs, as judged by immunoblotting, flow cytometry and Q-RT-PCR using ACE2-specific antibodies and primer/probes, respectively (**Supplemental Fig. 3 A-C**). In conclusion, freshly isolated, unstimulated PBMCs seem to be devoid of ACE2 expression. Furthermore, they appear to be non-susceptible and non-permissive to infection with either SARS-related coronavirus, at least *ex vivo*. However, the continuous presence of viral RNA associated to cells and in the culture supernatant suggests that virus particles attach to and/or internalize into PBMCs in an ACE2-independent manner and remain cell-associated for up to several days.

### Exposure of PBMCs to SARS-CoV-2, but not SARS-CoV, triggers a JAK/STAT-dependent cell-intrinsic innate immune response

To identify potential cell-intrinsic innate immune responses to SARS-CoV and SARS-CoV-2 exposure, we analyzed *IFIT1* and *IL6* mRNA expression over time (**Fig. 2A-B**). We selected *IFIT1* and *IL*6 as prototypic target genes that are transcribed by IRF3 and NF-κB, respectively (Honda and Taniguchi, 2006). In contrast to SARS-CoV-inoculated cells, SARS-CoV-2-exposed cells displayed Ruxolitinib-sensitive, significantly upregulated *IFIT1* mRNA expression at 16, 24 and 48 hours post-inoculation (**Fig. 2A**). Inhibition of potential low-level SARS-CoV-2 RNA replication through treatment of cells with Remdesivir, and heat-inactivation of the SARS-CoV-2 stock inoculum did not prevent induction of *IFIT1* mRNA expression (**Supplemental Fig. 4**), corroborating the idea that the latter is triggered by exposure to virions, but not by productive infection. In contrast, *IL6* expression was barely induced after exposure to SARS-CoV and SARS-CoV-2 (**Fig. 2B**). We next analyzed if type I IFN expression preceded *IFIT1* mRNA expression in SARS-CoV-2-exposed PBMCs. Despite a slight trend for elevated *IFNA1* and *IFNB1* mRNA expression at 16 hours, levels failed to reach significant upregulation at four, 16 and 24 hours, when compared to mock-exposed cultures (**Fig. 2C**). Together, SARS-CoV-2 exposure specifically triggered IRF3-induced *IFIT1*, but not NF-κB-transcribed *IL-6* gene expression. These results suggest that, although both SARS-related coronaviruses failed to establish a productive infection in PBMCs, SARS-CoV-2 appears to induce cell-intrinsic, JAK/STAT-dependent responses in several cell types comprised in PBMCs.

**Figure 2.**
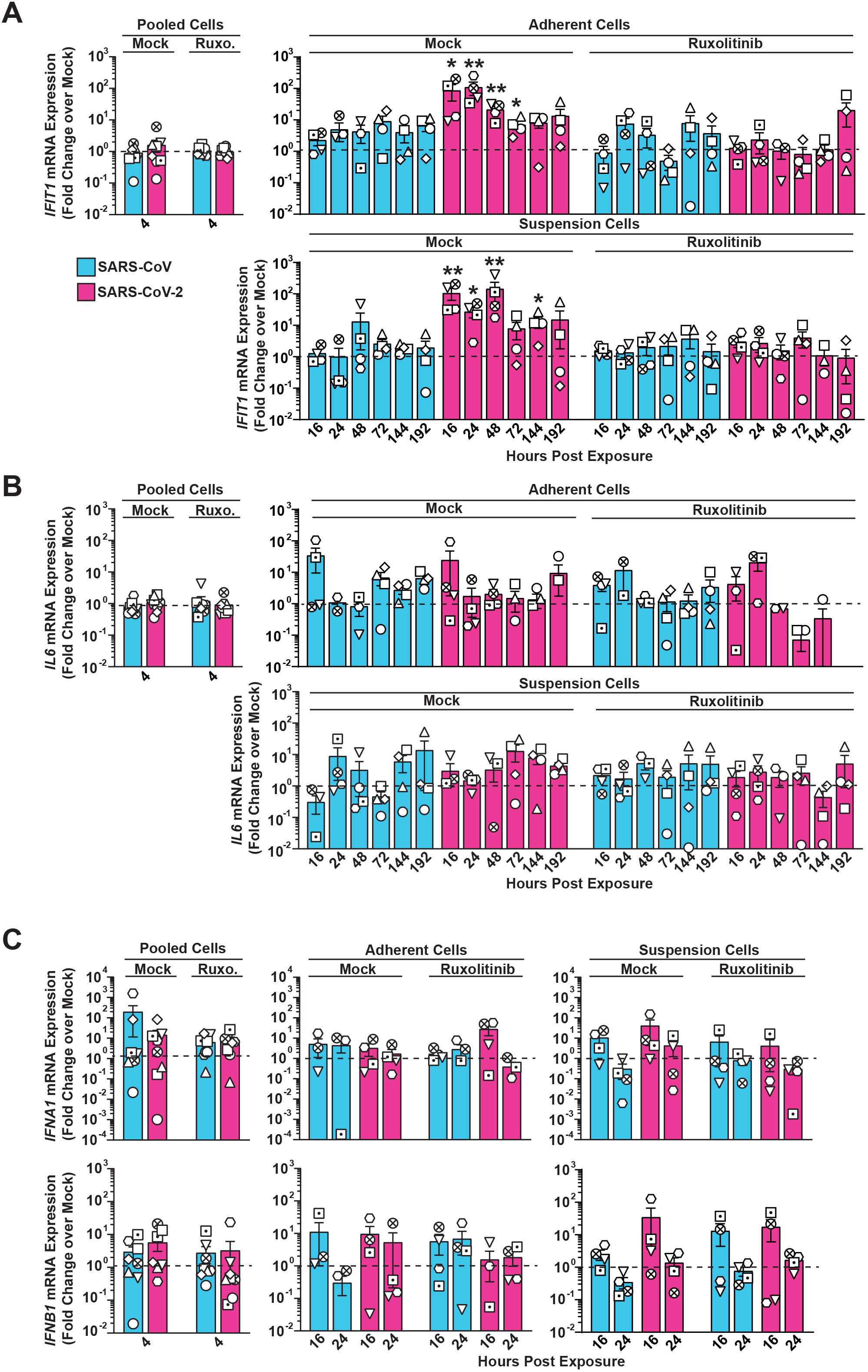
Exposure of PBMCs to SARS-CoV-2, but not SARS-CoV, triggers a JAK/STAT- dependent cell-intrinsic innate response. RNA extracted from Ruxolitinib-treated or mock-treated, and SARS-CoV-, SARS-CoV-2- or mock-exposed PBMCs was analyzed for: (**A**) *IFIT1*, (**B**) *IL-6*, (**C**) *IFNA1* and *IFNB1* mRNA expression by Q-RT-PCR at indicated time points. Suspension and adherent cell fractions were analyzed separately, except at the four hours time point. Values were normalized to cellular *RNASEP* expression and are shown as fold change over mock-inoculated conditions. The dotted line indicates the expression level of mock-inoculated cell cultures and is set to 1. Data were generated in four individual experiments using PBMCs from four individual donors represented by different symbols. n.d. = not detectable.

### SARS-CoV-2 exposure causes transcriptional changes in most cell types

To explore cell-intrinsic responses in individual cell types, we performed single-cell RNA-sequencing of PBMCs exposed to SARS-CoV and SARS-CoV-2, respectively. We identified the five major cell types, namely B-cells, CD4^+^ and CD8^+^ T-cells, NK cells and monocytes (**Fig. 3A**) based on the expression of discriminatory marker mRNAs (see methods). Separated based on experimental condition, PBMCs of both donors shared a similar relative cell type distribution (**Fig. 3B**) and similar cell type-specific transcriptional profile (**Supplemental Fig. 5**), and data of both donors were merged for the following analyses. In line with our bulk analyses (**Supplemental Fig. 3A-C**), *ACE2* mRNA was undetectable (**Supplemental Fig. 3E**), as was *TMPRSS2* mRNA. In contrast, the protease-encoding *FURIN*, *BSG* and *NRP1* mRNAs were expressed in all cell types, and most abundantly in monocytes (**Supplemental Fig. 3D-E**). Graphical mapping indicated transcriptomic changes within individual cell types for SARS-CoV-2-, but not for SARS-CoV-exposed cultures, compared to mock-inoculated cells (**Fig. 3C**). Notably, SARS-CoV-2 monocytes clustered separately from the other conditions in the UMAP despite library batch correction, implying a pronouncedly altered transcriptome. The T- and NK cell clusters slightly and partially shifted, indicating a change in their transcriptional profile (**Fig. 3C**). The relative abundance of T-cells and monocytes in SARS-CoV-2-exposed cells as compared to mock-exposed PBMCs remained constant, as judged by flow cytometric analysis (**Supplemental Fig. 6**). Together, this analysis revealed that transcriptomic changes occurred in most cell types upon SARS-CoV-2 exposure, particularly in the monocytic fraction.

**Figure 3.**
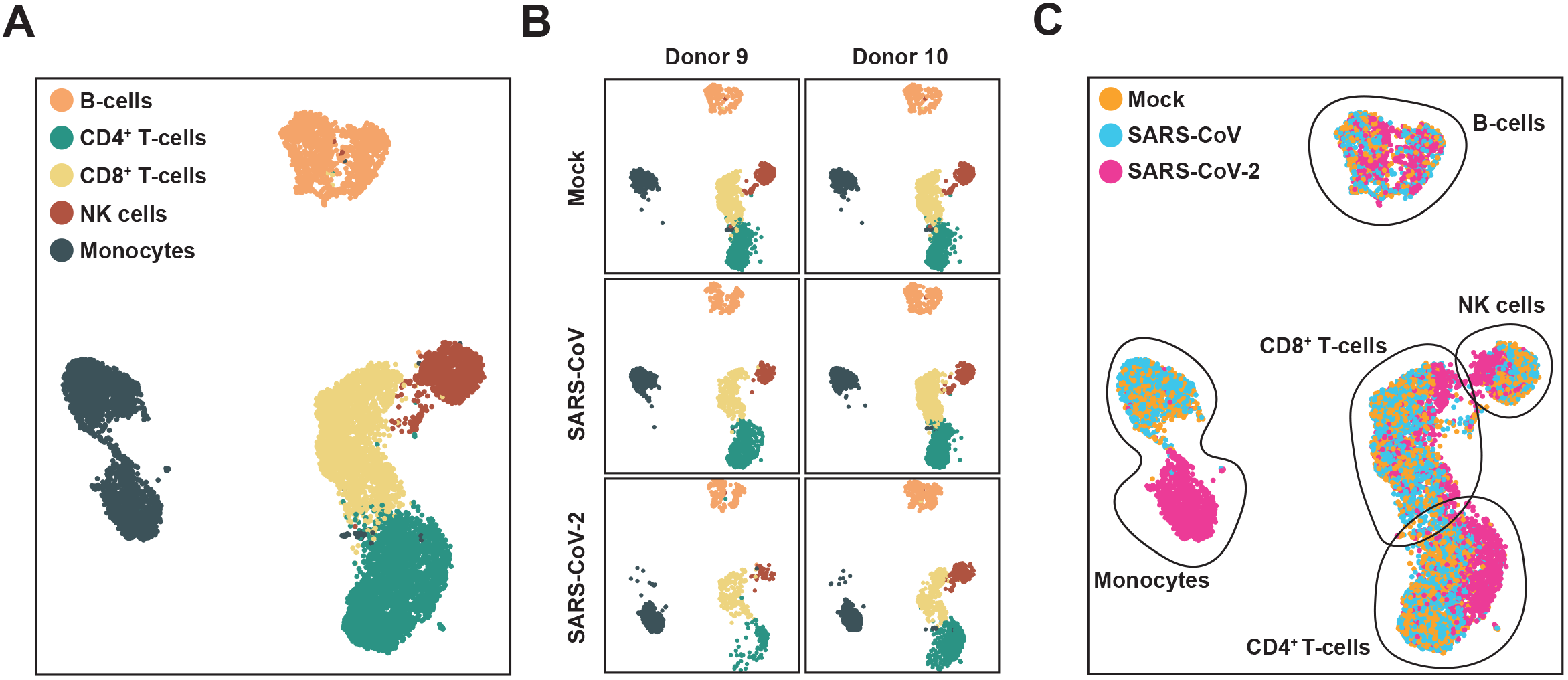
SARS-CoV-2 exposure causes transcriptional changes in most cell types. PBMCs isolated from two donors were exposed to SARS-CoV, SARS-CoV-2 or mock-exposed, and analyzed by scRNA-sequencing 24 hours post-exposure. (**A**) UMAP displaying all identified cell types, (**B**) UMAP indicating the data obtained from the PBMCs of the two donors, (**C**) Identified cell types according to condition.

### Exposure to SARS-CoV-2 induces a global innate immunity-related gene profile in PBMCs with cell type-specific signatures

We next investigated in more detail the cell type-specific response to SARS-CoV-2. Based on differentially expressed genes (DEGs) between mock-, SARS-CoV- and SARS-CoV-2-exposed PBMCs, a pseudotime cell trajectory analysis for all cell types was performed (**Fig. 4A**). For all five major cell types, cells inoculated with SARS-CoV-2 developed towards a separate cell fate and branched off from mock-exposed and SARS-CoV-exposed cells, which, conversely, shared a common trajectory. Interestingly, B-cell analysis resulted in four branching points, from which only two (#1 and #3) were specific for SARS-CoV-2-exposed cells, suggesting a high transcriptional heterogeneity of B-cells independently of virus exposure. Though progression through pseudotime resulted in a distinct and highly pronounced trajectory of all SARS-CoV-2-exposed cell types, this effect was most pronounced in monocytes (**Fig. 4A**). Analysis of expression of specific genes, including *ISG15* and *IFIT1,* confirmed that in general, all cell types contributed to gene expression changes upon SARS-CoV-2 challenge, and monocytes displayed the most pronounced elevation of expression of both genes (**Fig. 4B**). Identification of DEGs in mock-exposed compared to SARS-CoV-2-inoculated PBMCs revealed a significant upregulation of gene expression in all cell types, especially in monocytes (**Fig. 4C**). Interestingly, the majority of DEGs were identified as known ISGs (defined by the interferome database; colored in green (interferome.org; v2.01)). Scoring the individual cell types and conditions by their expression of an IFN-signaling module revealed a SARS-CoV-2-specific upregulated expression in all cell types, though this was most prominent in monocytes (**Fig. 4D**). Moreover, IFN module scores were colinear with Pseudotime scores along the SARS-CoV-2 trajectory, supporting the notion that SARS-CoV-2 exposure induces a development of PBMCs towards an antiviral phenotype. Increase of expression of several ISGs, including *ISG15, IFIT1, IFITM3, DDX58, IFIH, LY6E, MX2, IFI6, GBP1, BST2*, was detectable predominantly, but not exclusively, in monocytes (**Fig. 4E**), supporting the hypothesis that monocytes play a key role in the induction of cell-intrinsic innate immune response to SARS-CoV-2 stimulation. In line with our previous findings (**Fig. 2**), SARS-CoV-2- and SARS-CoV-exposed cells scored virtually negative for expression of various cytokines, including *IL6* (**Fig. 4E**) and *IFN* mRNAs (**Supplemental Fig. 7**), although they express IFN receptors (**Supplemental Fig. 7**). In conclusion, these data reveal a strong induction of cell-intrinsic innate immunity in SARS-CoV-2-exposed PBMCs that manifests predominantly in monocytes.

**Figure 4.**
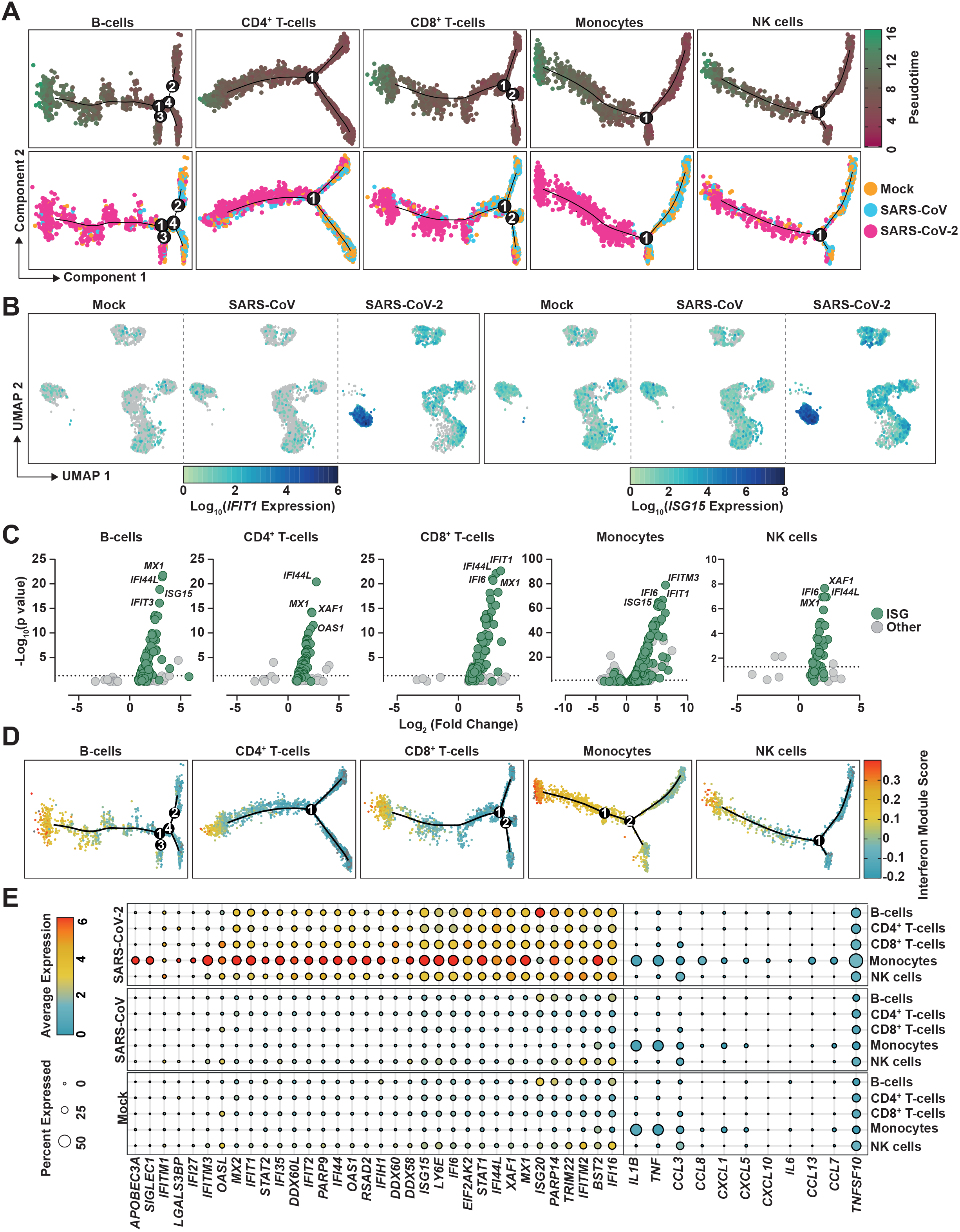
Exposure to SARS-CoV-2 induces a global innate immunity-related gene profile in PBMCs with cell type-specific signatures. (**A**) Pseudotime cell trajectory analysis and GSEA analysis using genes differentially regulated between mock-, SARS-CoV- and SARS-CoV-2-challenged conditions for indicated cell types. (**B**) Representative UMAPs showing *IFIT1* and *ISG15* mRNA expression in the indicated conditions. (**C**) Volcano plot of all DEGs in SARS-CoV-2-exposed cells compared to mock-exposed cells in the indicated cell types. Known ISGs were colored in green based on their presence in the interferome database (http://www.interferome.org/; v2.01). (**D**) Cell trajectory maps of indicated cell types with cells colored by expression of the genes in an IFN-module gene set. (**E**) Dot plot depicting expression of selected ISGs and cytokines. Expression levels are color-coded, the percentage of cells expressing the respective gene is coded by symbol size.

### Transcriptome differences in viral RNA-positive and bystander monocytes

Next, we aimed at identifying viral RNA-positive cells and their specific transcriptional profile that we hypothesized to differ from cells without detectable viral RNA of the identical culture. SARS-CoV-2 RNA was detectable in all cell types, but predominantly in monocytes (**Fig. 5A**). Identified viral reads were distributed over the viral genome sequence, with a high over-representation of the 3’ RNA sequences that all subgenomic and genomic viral RNA have in common, corresponding to the 3’ part of the N-coding sequence and polyA tail (**Fig. 5B**). Specifically, in SARS-CoV- and SARS-CoV-2-exposed PBMC cultures, we identified 99 (2.13%) and 212 (2.88%) viral RNA-positive cells, respectively (**Fig. 5C**). Among those, we identified 56 (7.8%) and 173 (15.3%) viral RNA-positive monocytes among all monocytes, respectively. First of all, no statistically significant differences in expression of individual genes of RNA-positive and RNA-negative monocytes were identified. However, the IFN module score (**Fig. 4**) was slightly, but statistically highly significantly elevated in SARS-CoV-2-exposed monocytes with undetectable viral RNA (**Fig. 5D-E**). Specifically, within the 94 genes that were expressed marginally more abundantly in cells lacking detectable SARS-CoV-2 RNA, 18 represented ISGs, including *ISG15*, *IFITM2*, *IFITM3*, *IFI27* and *HLA* genes tended to be upregulated in viral RNA-negative bystander cells. Importantly, the presence of viral RNA did not specifically associate with expression of *BSG*/*CD147* and *NRP1*, and *ACE2* and *TMPRSS2* expression was undetectable, suggesting that particles internalize in a manner that is independent of these confirmed or proposed receptors, respectively. In SARS-CoV-2 RNA-positive cells as compared to SARS-CoV-2 RNA-negative cells of the identical cultures, among others, *CD163* reads tended to be slightly more abundant. Expression of the hemoglobin-haptoglobin scavenger receptor CD163 has been associated with regulation of inflammation (Kowal et al. 2011) and has interestingly been linked to immunological changes in monocytes and monocyte-derived macrophages from SARS-CoV-2-infected individuals (Gómez-Rial et al. 2020; Trombetta et al. 2021; Wendisch et al. 2021). Looking specifically at the *CD163*^HIGH^ monocyte population, we found that it displayed high expression levels of genes with profibrotic functions, in line with, including *VCAN*, *LGMN*, *MERTK*, *TFGB1*, *MRC1*, *TGFBI* and *MMP9* and enhanced expression of cytokines including *CCL2*, *CXCL8* or *IL1B* and the cytokine receptor *CCR5* (**Suppl. Fig. 8**). Furthermore, SARS-CoV-2 RNA-positive cells displayed a preferential upregulation of genes implicated in migration and integrin binding (*FN1*, *PPBP*, *THBS1*) as well as differentiation, including *FABP5* and *LGMN*. Together, cells that internalized SARS-CoV-2 particles exhibit a slightly distinct gene expression profile characterized by a consistent reduction of antiviral ISGs and an upregulation of pro-fibrotic genes as opposed to bystander cells with undetectable viral RNA.

**Figure 5.**
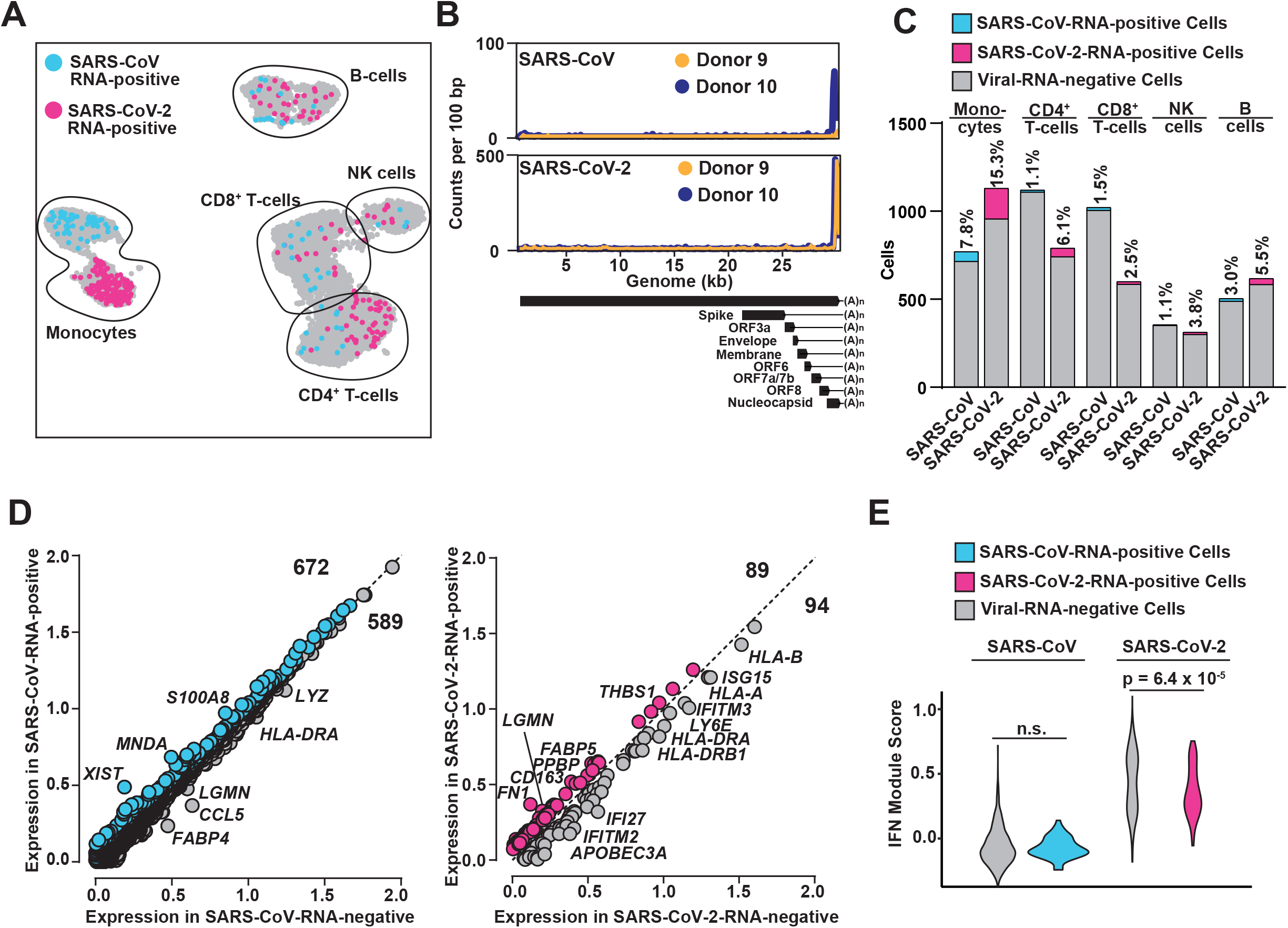
Viral RNA-positive monocytes trend toward downregulation of ISGs and upregulation of fibrosis-associated genes. (**A**) UMAP highlighting cells in which transcripts of either SARS-CoV RNA (blue) or SARS-CoV-2 RNA (magenta) were identified. (**B**) Virus-specific reads were aligned to the SARS-CoV or SARS-CoV-2 genome. Coverage of the genome is shown in counts per 100 bp. (**C**) Bar graph showing the absolute number (bars) and relative percentage of cells that were identified as virus RNA-positive in SARS-CoV- and SARS-CoV-2-inoculated cultures, respectively. (**D**) Plot of Log_10_ average expression of genes showing a Log_2_(fold change) >0.2 in viral RNA-positive versus viral RNA-negative monocytes from the SARS-CoV-(left panel) and SARS-CoV-2- (right panel) inoculated PBMCs with genes showing the highest expression fold change between both conditions. **(E)** IFN Module Score of viral-RNA-negative (grey), SARS-CoV-RNA-positive (blue) and SARS-CoV-2-RNA-positive (magenta) monocytes. Statistical significance was tested using a Wilcoxon rank sum test with continuity correction.

## DISCUSSION

In this study we characterized the response of peripheral immune cells, at the cell type level and at individual cellś level, to *ex vivo* SARS-CoV-2 exposure as compared to SARS-CoV. While *ex vivo* experiments inherently do not recapitulate systemic immune cell interactions and lack the context of complex tissueś and organś interplay and communication, they uniquely allow the side-by-side comparison of two genetically closely related, but functionally different viruses under standardized conditions. Furthermore, they allow assessing the direct consequence of virus exposure on individual cell types.

Our results indicate that SARS-CoV-2 and SARS-CoV share inability to detectably infect PBMCs. Previous studies with SARS-CoV and MERS-CoV yielded partially conflicting results regarding the susceptibility of human PBMCs to infection. *Ex vivo*, one publication reported absence of SARS-CoV replication in PBMCs (Castilletti et al. 2005), while another work suggested susceptibility and permissiveness of PBMCs to SARS-CoV infection with a high inter-donor variability (L. F. P. Ng et al. 2004). *In vivo*, *in situ* hybridization and electron microscopy analyses reported presence of SARS-CoV material in lymphocytes and monocytes derived from infected patients (Gu et al. 2005). MERS-CoV was suggested to efficiently replicate in *ex vivo*-infected monocytes (Chu et al. 2014), but to only abortively infect human T-cells (Chu et al. 2016). Of note, the confirmed receptor for SARS-CoV-2 cell entry (Hoffmann et al. 2020) has been reported to be virtually absent in PBMCs (Song et al. 2020; Xu et al. 2020; Zou et al. 2020; Xiong et al. 2020), a finding that is in line with our own inability to detect *ACE2* mRNA and ACE2 protein expression in PBMCs by various methods. Therefore, we hypothesize that virus particles attach and/or internalize in an ACE2-independent manner, resulting in viral RNA associated to and/or internalized into cells. Given that receptor-independent phagocytosis is a hallmark of monocytes, our observation that the majority of the viral reads were retrieved in monocytic cells underlines this idea. Furthermore, as SARS-CoV ORF7a is a virion-associated protein (Cheng Huang et al. 2006) and SARS-CoV-2 ORF7a was reported to efficiently interact with PBMC-derived monocytes (Zhou et al. 2021), ORF7a may contribute to attachment to monocytes. Interestingly, the binding capability of SARS-CoV ORF7a protein was reported to be significantly weaker as compared to SARS-CoV-2 ORF7a (Zhou et al. 2021), which is consistent with the observed two-fold reduced proportion of virus RNA-positive monocytes in SARS-CoV-exposed PBMCs as compared to SARS-CoV-2.

*In vivo*, a multitude of cytokines, including IL-1β, IL-1RA, IL-7, IL-8, IL-9, IL-10, CXCL10, IFN-γ and TNF-α are upregulated in the plasma of COVID-19 patients, especially in cases with severe outcome (Chaolin Huang et al. 2020). In contrast, mild COVID-19 associates with effective type I IFN responses, including expression of type I IFNs themselves and IFN-stimulated genes, that are probably essential to clear the virus infection and orchestrate adaptive immunity accordingly. To date, it remains largely unclear which cell populations are the drivers of these individual responses. Productively infected epithelial cells in the respiratory tract may initiate some of these responses directly; alternatively, or in addition, immune cells may be stimulated by signals released by productively infected cells or by virions and/or viral proteins directly. Studies on the consequence of the physical interaction of SARS-CoV-2 with infection-refractory primary immune cells, as opposed to susceptible cell types in the respiratory tract, are largely missing. Of note, cytokines levels and composition differ in serum and bronchoalveolar lavage fluid of patients with COVID-19 (Xiong et al. 2020), suggesting that productively infected epithelial tissue in the respiratory tract and nonsusceptible peripheral immune cells initiate different cytokine responses. Pro-inflammatory monocytes that infiltrate the lung have been proposed to represent major cytokine producers in the lung microenvironment (Liao et al. 2020). In line with this idea, SARS-CoV-2-susceptible infected cell lines and primary cells (Blanco-Melo et al. 2020) display imbalanced host responses, with strong cytokine and ablated ISG responses, when compared to other respiratory virus infections. Also, studies performed in the SARS-CoV-2 Syrian hamster model uncovered an early and strong cytokine response in the myeloid compartment of the lung (Nouailles et al. 2021). Here, our data provide first insight on the response of refractory PBMCs upon exposure to virus particles in the absence of co-stimulating infected cell types. The lack of expression of pro-inflammatory cytokines, including IL-6, TNFa and IL-1 in SARS-CoV-2-exposed PBMCs is in line with the idea that these cytokines are mainly derived from the respiratory tract representing the site of productive infection and it may partially explain the absence of lymphocyte depletion in our experimental setting that is observed *in vivo* (Chaolin Huang et al. 2020; Qin et al. 2020; D. Wang et al. 2020). In our *ex vivo* PBMC setting, which is devoid of productive infection, SARS-CoV-2, but not SARS-CoV particles induced innate immune responses in the absence of coculture with infected epithelial cells, indicating that direct exposure with virions can trigger responses in PBMCs.

Immune responses were initiated in different cell types with a focus on monocytes and were characterized by ample induction of expression of *IFIT1* and several other ISGs, as opposed to pro-inflammatory cytokines, including *IL6* mRNA expression. Our data suggest that this response may be triggered, at least to a certain extent, in a virus replication-independent manner. Despite our failure to detect IFN gene expression at the time points investigated, the Ruxolitinib-sensitive induction of *IFIT1* expression strongly suggests an underlying IFN signaling-dependent mechanism. Comparison of cells with and without detectable SARS-CoV-2 RNA revealed quantitative differences regarding gene expression. Genes associated to fibrosis, migration and integrin binding were mildly upregulated in cells with detectable viral RNA when compared to bystander cells, defined as cells of the SARS-CoV-2-exposed cell culture which lacked detectable viral reads. Interestingly, monocytes developing profibrotic functions have recently been established in the context of COVID-19 *in vivo* (Wendisch et al. 2021). Bystander cells displayed enhanced ISG expression, suggesting either a more efficient and more probable internalization into cells with a low ISG profile or, alternatively, delivery of a SARS-CoV-2-specific sensing antagonist in the context of efficient particle internalization. Multiple SARS-related coronaviruses-encoded IFN antagonists, including structural components of the incoming virion that do not require productive infection for being expressed and functional, dampen innate immune responses when ectopically expressed, including membrane and nucleocapsid proteins (Lei et al. 2020). In addition, virion components including ORF3 and ORF6 (Bai et al. 2021; Cheng Huang, Peters, and Makino 2007; Ito et al. 2005) have type I IFN evasion properties (Li et al. 2020; Schroeder et al. 2021; Lei et al. 2020)). Interestingly, among those, ORF6 from SARS-CoV-2 was described to be inferior in counteracting phospho-IRF3 nuclear translocation in infected cells, compared to SARS-CoV ORF6, resulting in higher ISG induction (Schroeder et al. 2021). Therefore, incoming viral RNA sensing may be less efficiently prevented by SARS-CoV-2 ORF6 as compared to SARS-CoV ORF6. Finally, the large absence of a detectable ISG expression profile in SARS-CoV-exposed PBMCs is consistent with a previous report analyzing abortively infected monocyte-derived macrophages (Cheung et al. 2005).

Together, our study provides analysis of gene expression in *ex vivo-*exposed PBMCs at the cell type and individual celĺs level. Our data suggest that direct stimulation of monocytes through physical contact to SARS-CoV-2 particles is followed by strong ISG induction, despite absence of detectable productive infection.

## METHODS

### Cell Lines and Primary Cells

Vero E6 (ATCC CRL-1586) cells, Calu-3 (ATCC HTB-55) cells and HEK293T (ATCC CRL-3216) cells were cultivated in Dulbecco’s modified Eagle’s medium (DMEM) supplemented with 10% heat-inactivated fetal calf serum, 1% non-essential amino acids (Thermo Fisher Scientific) and 1% sodium pyruvate (Thermo Fisher Scientific) in a 5% CO_2_ atmosphere at 37°C. Cell lines were routinely monitored for absence of mycoplasma and paramyxovirus simian virus 5.

Withdrawal of blood samples from healthy humans and cell isolation were conducted with approval of the local ethics committee (Ethical review committee of Charité Berlin, votes EA4/166/19 and EA4/167/19). Human PBMCs were isolated from buffy coats by Ficoll-Hypaque centrifugation. PBMCs were cultured at 2 × 10^6^/ml in RPMI 1640 containing 10% heat-inactivated fetal calf serum (Sigma-Aldrich), 1% penicillin-streptomycin (Thermo Fisher Scientific) and 2 mM L-glutamine (Thermo Fisher Scientific).

### Viruses

SARS-CoV isolate HKU-39849 (accession no. JQ316196.1, (Zeng et al. 2003; van den Worm et al. 2012)) and the SARS-CoV-2 BetaCoV/Munich/ChVir984/2020 isolate (B.1 lineage, EPI_ISL_406862, (Wölfel et al. 2020)) were used.

Virus was grown on Vero E6 cells and concentrated using Vivaspin® 20 concentrators with a size exclusion of 100 kDa (Sartorius Stedim Biotech) in order to remove cytokines of lower molecular weight, including IFNs. Virus stocks were stored at -80°C, diluted in OptiPro serum-free medium supplemented with 0.5% gelatine and PBS. Titer was defined by plaque titration assay. Cells inoculated with culture supernatants from uninfected Vero cells mixed with OptiPro serum-free medium supplemented with 0.5% gelatine and PBS, served as mock-infected controls. All infection experiments were carried out under biosafety level three conditions with enhanced respiratory personal protection equipment.

### Plaque Titration Assay

The amount of infectious virus particles was determined via plaque titration assay. Vero E6 cells were plated at 3.5 x 10^5^ cell/ml in 24-well and infected with 200 µl of a serial dilution of virus-containing cell culture supernatant diluted in OptiPro serum-free medium. One hour after adsorption, supernatants were removed and cells overlaid with 2.4% Avicel (FMC BioPolymers) mixed 1:1 in 2xDMEM. Three days post-infection, the overlay was removed, cells were fixed in 6% formaldehyde and stained with a 0.2% crystal violet, 2% ethanol and 10% formaldehyde. Plaque forming units were determined from at least two dilutions for which distinct plaques were detectable.

### Virus Exposure of PBMCs

30 min prior to virus exposure, PBMCs were left mock-treated or treated with Ruxolitinib (10 µM) or Remdesivir (20 µM). Treatment was maintained for the duration of the entire experiment. Virus challenge occurred by inoculation of 0.4 x 10^6^ cells/ml in RPMI cell culture medium supplemented with 2% FCS. Four hours post-challenge, cells were centrifuged and supernatants were collected (referred to as inoculum). Cells were resuspended in RPMI cell culture medium supplemented with 10% FCS and plated at 0.4 x 10^6^ cell/1.5 ml in 12-wells. In addition, post-wash samples were collected. For further sampling, cell culture supernatant was centrifuged, supernatant was collected and mixed with OptiPro serum-free medium supplemented with 0.5% gelatine for titration on Vero E6 cell or mixed with RAV1 buffer for viral RNA extraction and stored at -80°C until sample processing. Suspension cells and adherent cells were lysed in Trizol reagents and subjected to total RNA extraction.

### Reagents and Inhibitors

Ruxolitinib was purchased from InvivoGen and used at 10 μM concentration. Remdesivir (Gilead Sciences) was kindly provided by the Department of Infectious Diseases and Respiratory Medicine, Charité - Universitätsmedizin Berlin.

### Quantitative RT-Q-PCR

Viral RNA was extracted from cell culture supernatants using the NucleoSpin RNA virus isolation kit (Macherey-Nagel) according to the manufacturer’s instructions. Total RNA extraction from cells and DNase treatment were performed with Direct-zol RNA extraction kit (Zymo Research). Viral genome equivalents were determined using a previously published assay specific for both SARS-CoV and SARS-CoV-2 E gene (Corman et al., 2020). Subgenomic E gene expression was analyzed using the same probe and reverse primer combined with a forward primer, which is located in the SARS-CoV-2 leader region (sgLead-CoV-F: CGA TCT CTT GTA GAT CTG TTC TC (Wölfel et al. 2020)).

To analyze human gene expression, extracted RNA extraction was subjected to cDNA synthesis (NEB, Invitrogen). Quantification of relative mRNA levels was performed with the LightCycler 480 Instrument II (Roche) using Taq-Man PCR technology. For human *IFIT1* and *IFNB1*, a premade primer-probe kit was used (Applied Biosystems, assay IDs: Hs01911452_s1; Hs01077958_s1; respectively). For human *ACE2* (ACE2-F: TGCCTATCC TTCCTATATCAGTCCAA, ACE2-R:GAGTACAGATTTGTCCAAAATCTAC, ACE2-P: 6-FAM/ATGCCTCCCTGCTCATTTGCTTGGT/IBFQ), *IL-6* (IL-6-F: GGATTCAATGAGGAGACTTGC, IL-6-R: CACAGCTCTGGCTTGTTCC, IL-6-P: 6-FAM/AATCATCAC/ZEN/TGGTCTTTTGGAGTTTGAGG/IBFQ) and *IFNA1* (IFNA1-F:GGGATGAGGACCTCCTAGACAA, IFNA1-R:CATCACACAGGCTTCCAAGTCA, IFNA1-P:6-FAM/TTCTGCACCGAACTCTACCAGCAGCTG/BHQ), customer-designed oligonucleotides were synthesized by Integrated DNA Technologies (IDT). Relative mRNA levels were determined using the ΔΔCt method using human *RNASEP* (Applied Biosystems) as internal reference. Data analysis was performed using LightCycler Software 4.1 (Roche).

### Immunoblotting

Cells were washed once with ice-cold PBS and lysed in 60 µl RIPA Lysis Buffer (Thermo Fisher Scientific) supplied with 1% protease inhibitor cocktail set III (Merck Chemicals) for 30 min at 4°C. Cell debris were pelleted for 10 min at 13000g and 4°C and the supernatant was transferred to a fresh tube. Protein concentration was determined with the Thermo Scientific’s Pierce™ BCA protein assay kit according to the manufacturer’s instructions. Protein lysates were mixed with 4 X NuPAGE LDS Sample Buffer (Invitrogen) supplied with 10% 2-mercaptoethanol (Roth) and inactivated for 10 min at 99°C. Proteins were separated by size on a 12% sodium dodecyl sulfate polyacrylamide gel and blotted onto a 0.2 µm PVDF membrane (Thermo Scientific) by semi-dry blotting (BioRad). Human ACE2 was detected using a polyclonal goat anti-human ACE2 antibody (1:500, R&D Systems), a horseradish peroxidase (HRP)-labeled donkey anti-goat antibody (1:5000, Dinova) and Super Signal West Femto Chemiluminescence Substrate (Thermo Fisher Scientific). As a loading control, samples were analyzed for ꞵ-Actin expression using a mouse anti-ꞵ-actin antibody (1:5000, Sigma Aldrich) and a HRP-labeled goat anti-mouse antibody (1:10000, Dianova).

### Single-Cell RNA-Seq

Single-Cell RNA-seq libraries were prepared with the 10x Genomics platform using the Chromium Next GEM Single Cell 3’ Reagent Kits v.3.1 following manufacturer’s instructions. Samples were multiplexed using TotalSeq-A Antibodies purchased from BioLegend (A0256, A0258 and A0259). Antibody staining and the subsequent library preparation were performed following manufacturer’s instructions. Quality control of the libraries were performed with the KAPA Library Quantification Kit and Agilent TapeStation. Libraries were sequenced on a HiSeq4000 using the following sequencing mode: read 1: 28 bp, read 2: 91-100 bp, Index i7: 8 bp. The libraries were sequenced to reach ∼20 000 reads per cell.

### Single-Cell RNA-Seq Data Analysis

BCL files from the sequencing protocol were processed using the Cell Ranger pipeline v 3.1.0 (10X Genomics) and further analysed using the Seurat v3.1.4 package (Butler et al. 2018) in R v3.6 (https://www.r-project.org/). Pre-processing of the data was performed using the recommended SCTransform procedure and the IntegrateData with PrepSCTIntegration workflows to eliminate batch effects. A comprehensive description of the code used in the analysis of data is available at https://github.com/GoffinetLab/SARS-CoV-2_PBMC-study. Cell types were identified based on marker gene expression (Schulte-Schrepping et al. 2020): B-cells (*CD3D*^-^, *MS4A1*^+^), CD4^+^ T-cells (*CD3D*^+^, *CD8A*^-^), CD8^+^ T-cells (*CD3D*^+^, *CD8A*^+^), NK cells (*CD3D*^-^, *CD8A*^-^, *NKG7*^+^, *GNLY*^+^), Monocytes *(CD3D*^-^, *CD14*^+^, *FCGR3A*^+^).

### Cell Trajectory Analysis

Cell trajectory analysis was performed using using the Monocle v2.14.0 package (Trapnell et al., 2014) according to the guidelines set out by the developers. Different cell types were subclustered and processed as mentioned above. A resolution parameter of 0.3 was used for clustering. DEGs between clusters were determined using Seurat’s FindAllMarkers function (Wilcoxon sum-rank test); of these, genes with a Bonferroni-corrected p-value of <0.05 were imputed as ordering genes to generate the minimum spanning tree using the DDRTree algorithm. Code available at https://github.com/GoffinetLab/SARS-CoV-2_PBMC-study.

### IFN Module Score

The IFN-signaling pathway gene set [R-HSA-913531] from the Reactome database (Bijay *et al*., 2020) was retrieved from the Molecular Signatures Database (MSigDB) (Liberzon *et al*., 2011). Cells were scored on their expression of these genes using the AddModuleScore function in Seurat, which is referred to as the IFN module score as the pathway includes genes canonically differentially regulated in response to interferon signaling.

### Flow Cytometry Analysis

PBS-washed cells were PFA-fixed and immunostained for individual surface protein expression using the following antibodies: Anti-CD3-FITC (#561807; BD Biosciences), anti-CD4-APC (#555349; BD Biosciences), anti-CD14-PE (#561707; BD Biosciences), anti-CD19-FITC (#21270193; ImmunoTools), anti-NRP1/CD304-APC-R700 (#566038, BD Biosciences), anti-PD-1/CD279-PE (#21272794; ImmunoTools) and anti-TIM-3/CD366-FITC (#345022; Biolegend). To determine ACE2 cell surface expression, cells were immunostained with a goat anti-human ACE2 antibody (#AF933, R&D Systems) followed by immunostaining with a secondary antibody donkey anti-goat Alexa Fluor 488 (#A-11055, Thermo Fisher). ACE2-positive HEK293T cells were generated by transduction of cells with retroviral vectors generated by transfection of HEK293T cells with MLV gag-pol (Bartosch, Dubuisson, and Cosset 2003), pCX4bsrACE2 (Kamitani et al. 2006) and pVSV-G (Stewart et al. 2003). A FACS Lyric device (Becton Dickinson, Franklin Lakes, New Jersey, USA) with BD Suite Software was used for analysis.

### Data Presentation and Statistical Analysis

If not stated otherwise, bars show the arithmetic mean of indicated amount of repetitions. Error bars indicate S.E.M. from the indicated amount of individual experiments. Statistical significance was calculated by performing Student’s t-test using GraphPad Prism. *P* values <0.05 were considered significant and marked accordingly: P<0.05 (*),P <0.01 (**) or P< 0.001; n.s. = not significant (≥0.05).

## Data availability

The raw sequencing datasets generated during this study will be made available at the NCBI Gene Expression Omnibus.

## Acknowledgments

We thank Julian Heinze for excellent technical support. We thank J. S. M. Peiris, University of Hong Kong, China for providing the SARS-SARS-CoV isolate HKU-39849. We are grateful for access to the BIH Core Facility Sequencing. This work was supported by funding from Berlin Institute of Health (BIH) to C.G. J.K. is supported by the Center of Infection Biology and Immunity (ZIBI) and Charité PhD Program. Part of this work was supported by Deutsche Forschungsgemeinschaft (DFG) (SFB TR84 to CD) and Bundesministerium für Bildung und Forschung (BMBF) through the projects RAPID-2 (01KI2006A) to CD. A.-E.S. thanks FOR-COVID (Bayerisches Staatsministerium für Wissenschaft und Kunst) and the Helmholtz Association for financial support.

## Author Contributions

J.K., K.F., D.N. and C.G. conceived and designed the experiments; J.K., K.F., J.J., A.R., L.B.D.J., C.S., M.S., and D.N. performed the experiments, J.K., K.F., D.P., C.F., L.B.D.J., D.N. and C.G. analyzed the data; C.D., A.E.S. and L.S. provided resources; D.N. and C.G. drafted the manuscript.; J.K., K.F., D.P., D.N. and C.G. reviewed and edited the manuscript; S.S., D.N. and C.G. supervised the research.

## Conflict of Interest

Technische Universität Berlin, Freie Universität Berlin and Charité - Universitätsmedizin have filed a patent application for siRNAs inhibiting SARS-CoV-2 replication with D.N. as co-author.

---

**Supplemental Figure 1.**
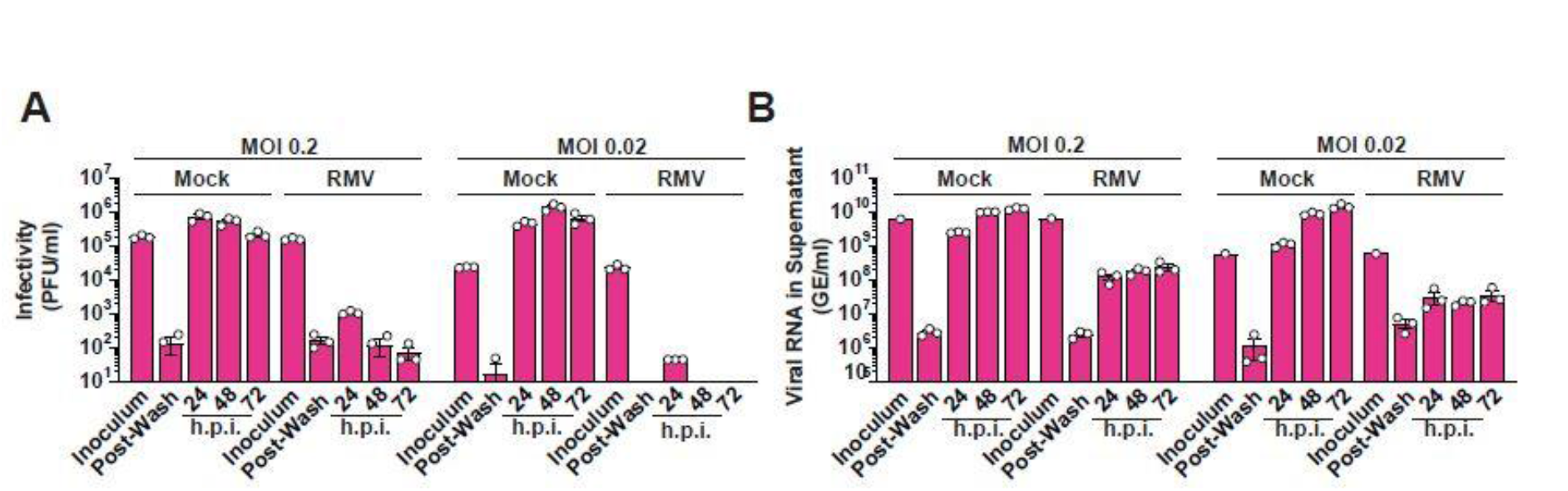
Remdesivir-sensitive SARS-CoV-2 infection of Vero E6 cells. Vero E6 cells were treated with Remdesivir (20 µM) or mock-treated and infected with SARS-CoV-2 at indicated MOIs. At indicated time points post-infection, (**A**) infectivity in supernatant was quantified by plaque titration assays and (**B**) viral RNA in supernatant was quantified by Q-RT-PCR. RMV = Remdesivir.

**Supplemental Figure 2.**
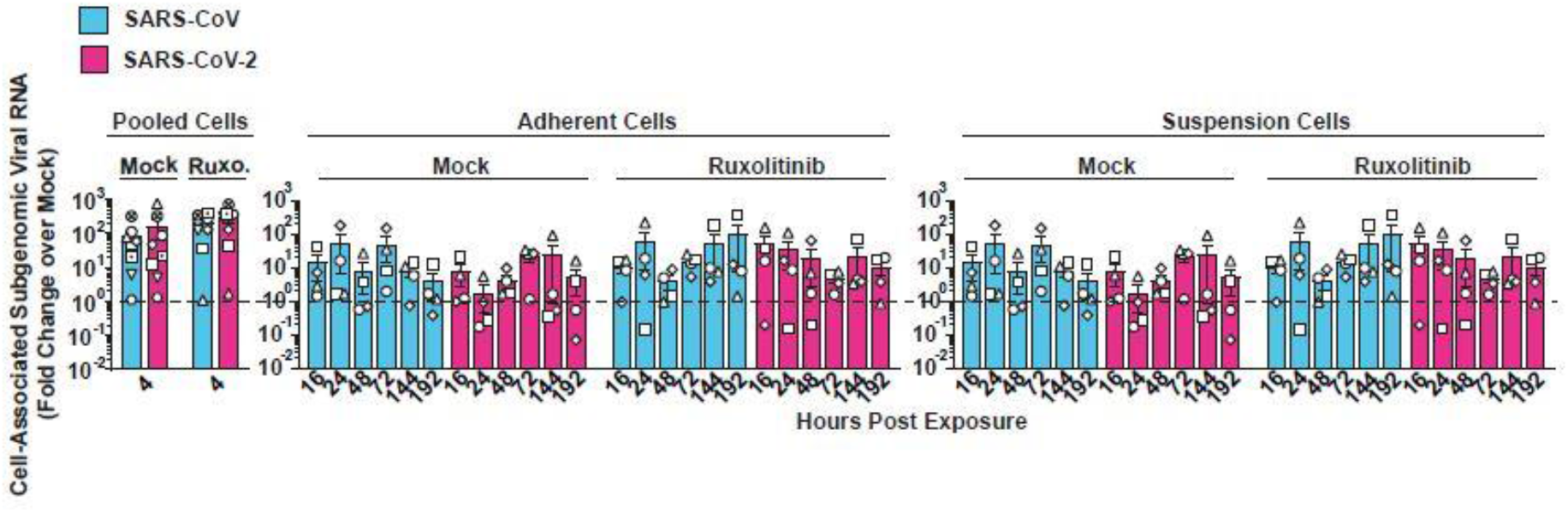
Absence of *de novo* production of subgenomic viral RNA in SARS-CoV- and SARS-CoV-2-inoculated PBMCs. PBMCs were treated with Ruxolitinib (10 µM) or mock-treated and challenged with SARS-CoV, SARS-CoV-2 or mock-challenged. Cell-associated subgenomic viral RNA (sgRNA encoding envelope gene) at indicated time points was quantified in suspension and adherent cells by Q-RT-PCR and normalized to mock-inoculated samples.

**Supplemental Figure 3.**
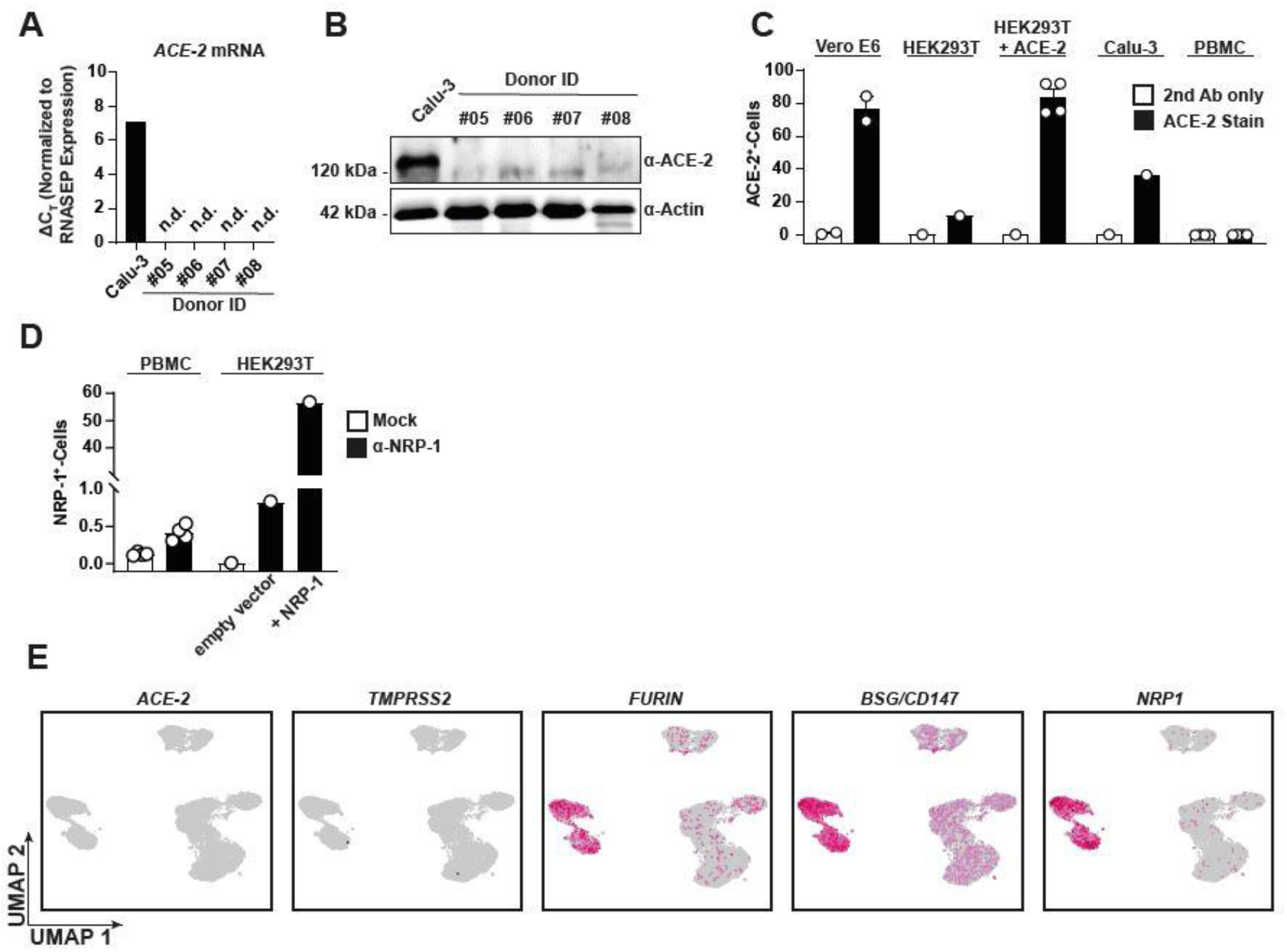
Cofactor expression profile in PBMCs. (**A-C**) ACE2 expression was analyzed in PBMC lysates from four healthy donors and Calu-3 cells by Q-RT-PCR of *ACE2* mRNA (**A**), anti-ACE2 immunoblotting (**B**), and anti-ACE2 immunostaining followed by flow cytometric analysis (**C**). (**D**) Surface NRP1 expression was quantified by flow cytometry in PBMCs and indicated HEK293T cells. (**E**) UMAPs showing expression of *ACE2*, *TMPRSS2*, *FURIN*, *BSG* and *NRP1* in all analyzed cells.

**Supplemental Figure 4.**
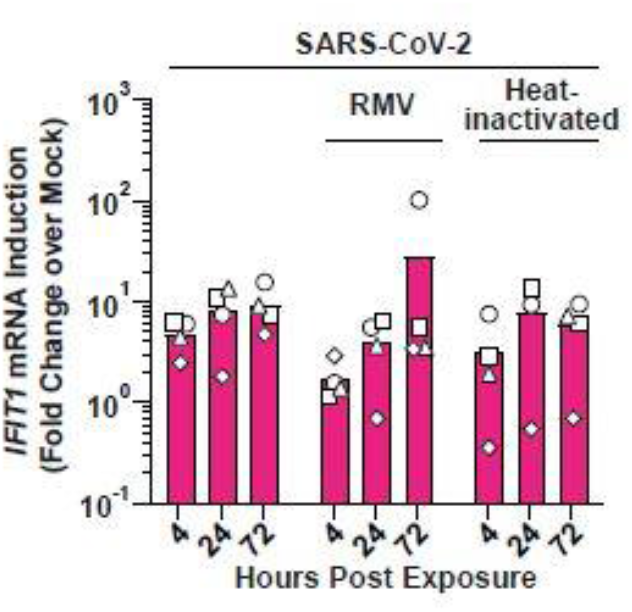
Exposure of PBMCs to SARS-CoV-2 induces *IFIT1* mRNA expression in the absence of productive infection. PBMCs were either mock-inoculated, pre-treated with Remdesivir or mock-treated, and challenged with SARS-CoV-2, or exposed to heat-inactivated SARS-CoV-2, and analyzed for *IFIT1* mRNA expression by Q-RT-PCR.

**Supplemental Figure 5.**
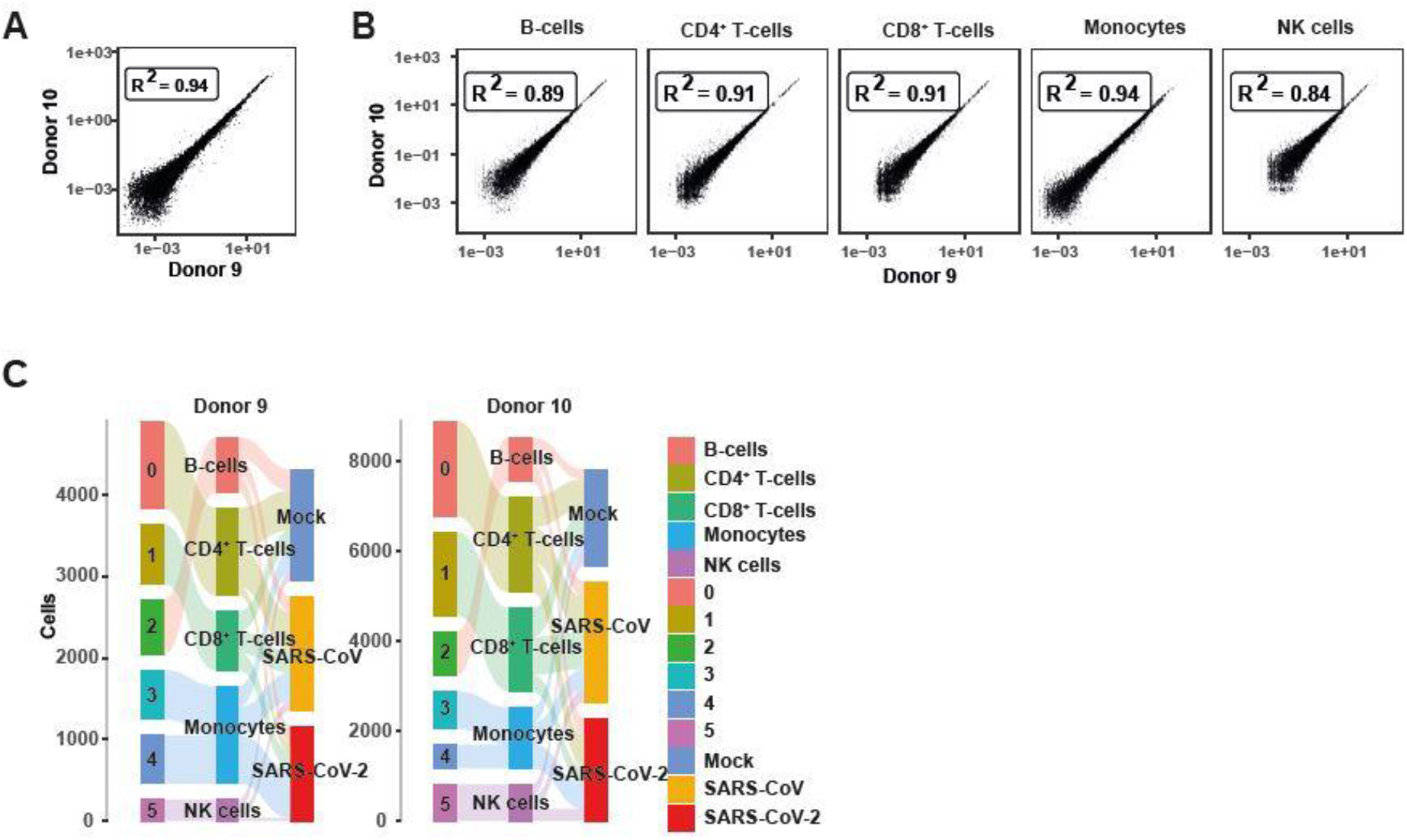
Quality control metrics of scRNA-seq data. Raw gene counts of (**A**) all identified genes in donor #9 and #10 or (**B**) separated by identified cell types. (**C**) Graph showing the cell number distribution of each sample in Seurat cluster (left), cell type annotations (middle) and treatments (right).

**Supplemental Figure 6.**
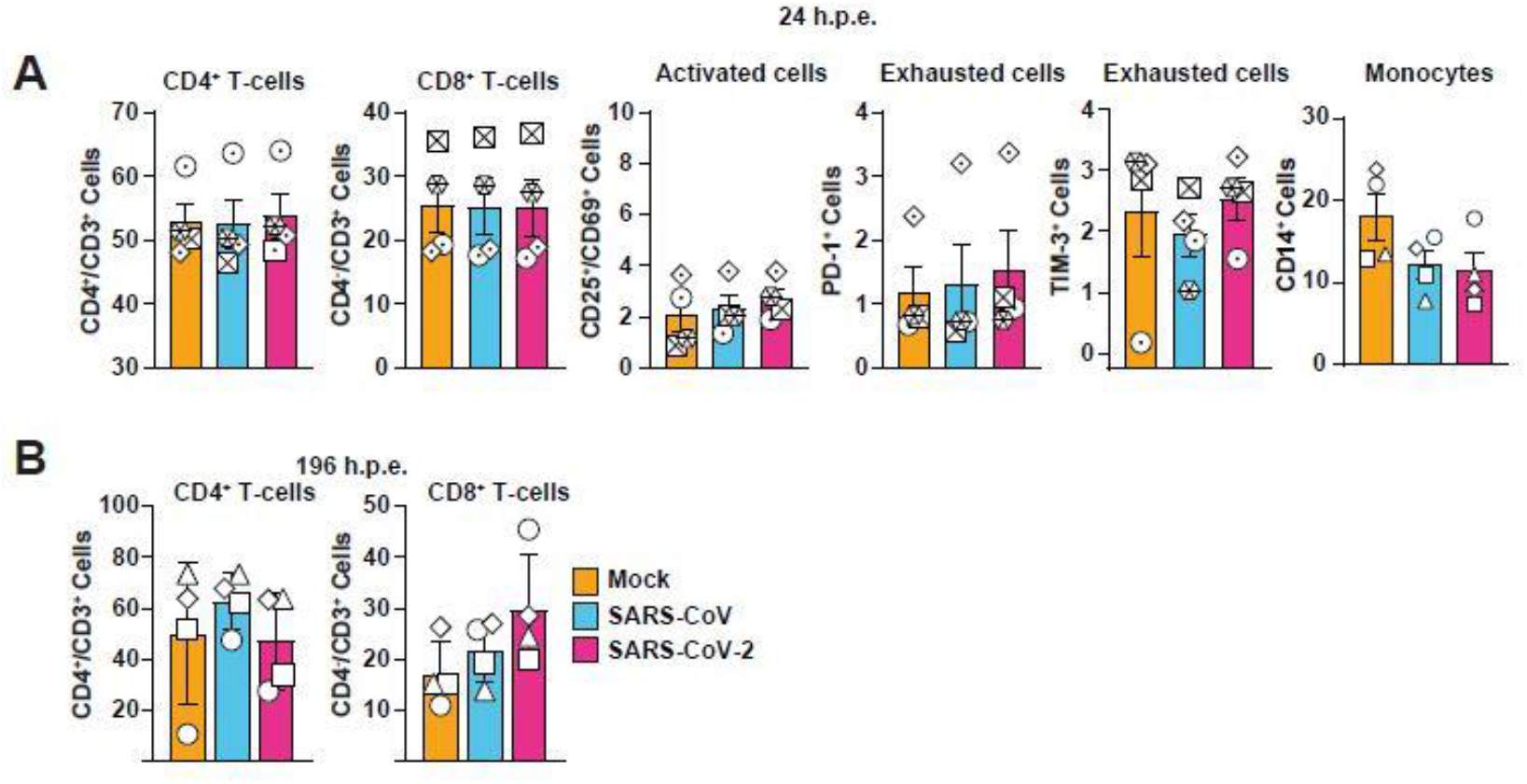
*Ex vivo* exposure with SARS-CoV or SARS-CoV-2 has no influence on T-cell activation and exhaustion. Surface immunostaining of SARS-CoV or SARS-CoV-2-exposed or mock-exposed PBMCs for T cell surface marker CD3, CD4, CD25/CD69, PD-1 and TIM-3 (**A**) 24 hours post-exposure or (**B**) 192 hours post-exposure. Symbols indicate cultures from individual donors, error bars indicate S.E.M. from four individual experiments.

**Supplemental Figure 7.**
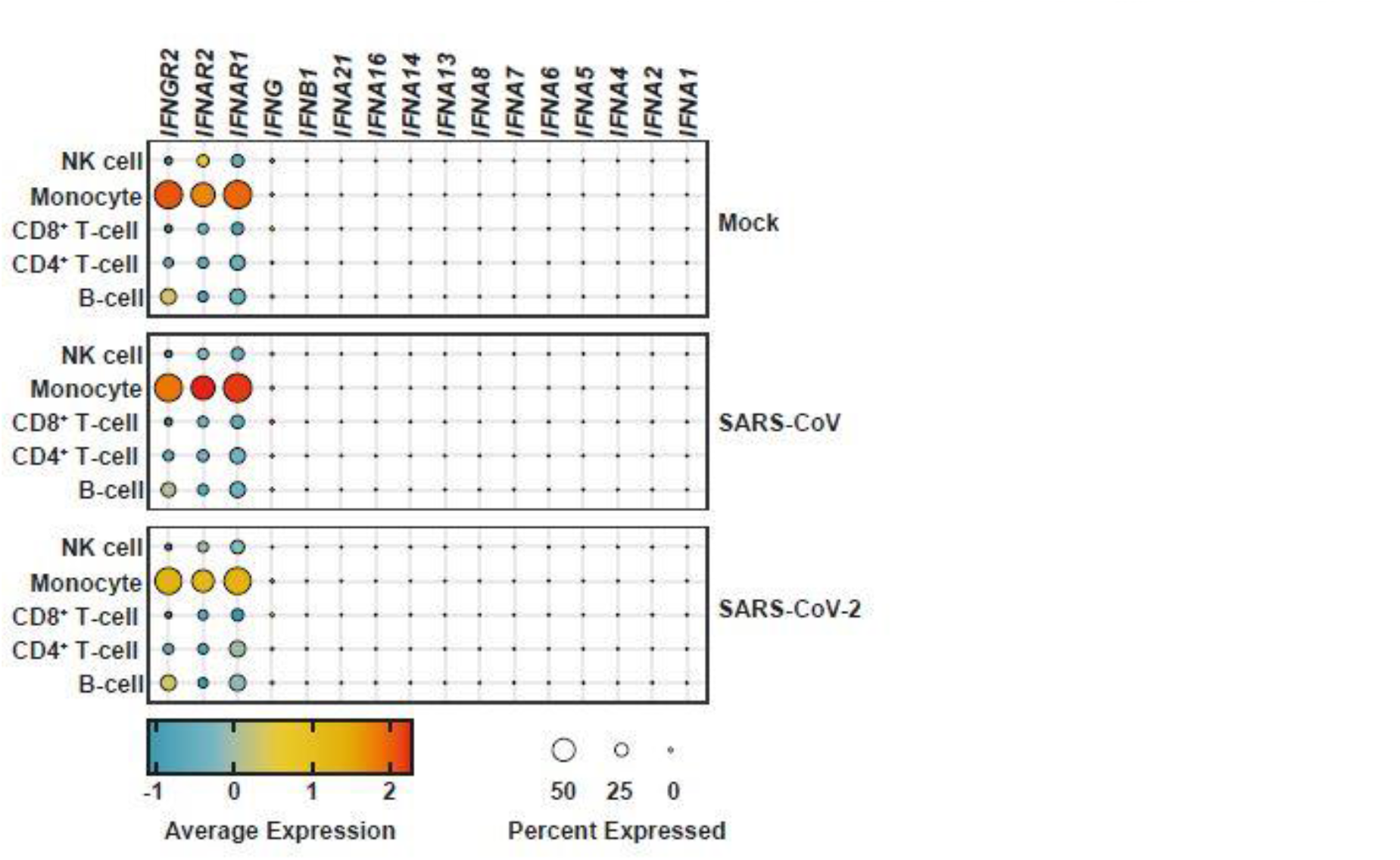
Expression of *IFNA*, *IFNB1, IFNG* genes and genes encoding for the IFN receptors. Dot plot showing the expression of the IFN-receptor encoding genes *IFNGR2*, *IFNAR1*, *IFNAR2* and individual *IFNA* subtypes, *IFNB1* or *IFNG* genes according to the treatment and cell types.

**Supplemental Figure 8.**
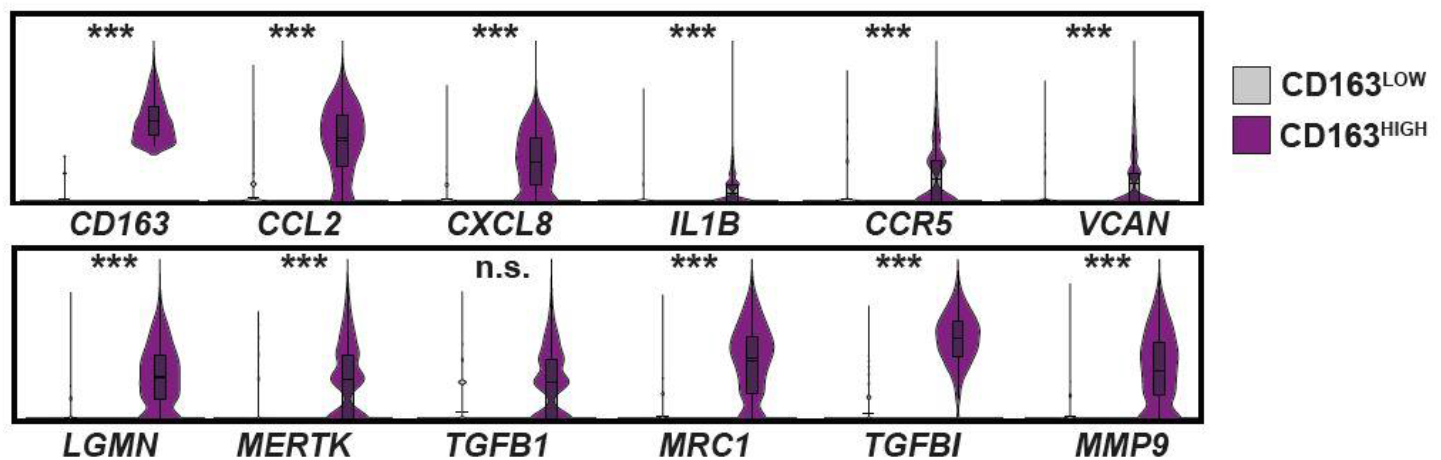
CD163^HIGH^ monocytes associate with marker genes specific for fibrosis. Distribution of indicated gene expression in CD163^HIGH^ and CD163^LOW^ monocytes. Cells were considered as CD163^HIGH^ with an Log_2_(CD163 average expression) >2. A total of 1520 CD163^HIGH^ and 20962 CD163^LOW^ cells were analyzed. *P* values <0.05 were considered significant and marked accordingly: P<0.05 (*), and P <0.01 very significant (**) or P< 0.001; n.s. = not significant (≥0.05).

## References

1. Anderson, Elizabeth M., Eileen C. Goodwin, Anurag Verma, Claudia P. Arevalo, Marcus J. Bolton, Madison E. Weirick, Sigrid Gouma, et al. 2021. “Seasonal Human Coronavirus Antibodies Are Boosted upon SARS-CoV-2 Infection but Not Associated with Protection.” Cell 184 (7): 1858–64.e10.

2. Andersson, Monique I., Carolina V. Arancibia-Carcamo, Kathryn Auckland, J. Kenneth Baillie, Eleanor Barnes, Tom Beneke, Sagida Bibi, et al. 2020. “SARS-CoV-2 RNA Detected in Blood Products from Patients with COVID-19 Is Not Associated with Infectious Virus.” Wellcome Open Research 5 (October): 181.

3. Arunachalam, Prabhu S., Florian Wimmers, Chris Ka Pun Mok, Ranawaka A. P. M. Perera, Madeleine Scott, Thomas Hagan, Natalia Sigal, et al. 2020. “Systems Biological Assessment of Immunity to Mild versus Severe COVID-19 Infection in Humans.” Science 369 (6508): 1210–20.

4. Bacher, Petra, Elisa Rosati, Daniela Esser, Gabriela Rios Martini, Carina Saggau, Esther Schiminsky, Justina Dargvainiene, et al. 2020. “Low-Avidity CD4+ T Cell Responses to SARS-CoV-2 in Unexposed Individuals and Humans with Severe COVID-19.” Immunity 53 (6): 1258–71.e5.

5. Bai, Zhihua, Ying Cao, Wenjun Liu, and Jing Li. 2021. “The SARS-CoV-2 Nucleocapsid Protein and Its Role in Viral Structure, Biological Functions, and a Potential Target for Drug or Vaccine Mitigation.” Viruses 13 (6). https://doi.org/10.3390/v13061115.

6. Bartosch, Birke, Jean Dubuisson, and François-Loïc Cosset. 2003. “Infectious Hepatitis C Virus Pseudo-Particles Containing Functional E1-E2 Envelope Protein Complexes.” The Journal of Experimental Medicine 197 (5): 633–42.

7. Bastard, P., L. B. Rosen, Q. Zhang, E. Michailidis, H. H. Hoffmann, Y. Zhang, K. Dorgham, et al. 2020. “Auto-Antibodies against Type I IFNs in Patients with Life-Threatening COVID-19.” Science. https://doi.org/10.1126/science.abd4585.

8. Blanco-Melo, Daniel, Benjamin E. Nilsson-Payant, Wen-Chun Liu, Skyler Uhl, Daisy Hoagland, Rasmus Møller, Tristan X. Jordan, et al. 2020. “Imbalanced Host Response to SARS-CoV-2 Drives Development of COVID-19.” Cell 181 (5): 1036–45.e9.

9. Bost, Pierre, Amir Giladi, Yang Liu, Yanis Bendjelal, Gang Xu, Eyal David, Ronnie Blecher-Gonen, et al. 2020. “Host-Viral Infection Maps Reveal Signatures of Severe COVID-19 Patients.” Cell 181 (7): 1475–88.e12.

10. Braun, Julian, Lucie Loyal, Marco Frentsch, Daniel Wendisch, Philipp Georg, Florian Kurth, Stefan Hippenstiel, et al. 2020. “SARS-CoV-2-Reactive T Cells in Healthy Donors and Patients with COVID-19.” Nature 587 (7833): 270–74.

11. Butler, Andrew, Paul Hoffman, Peter Smibert, Efthymia Papalexi, and Rahul Satija. 2018. “Integrating Single-Cell Transcriptomic Data across Different Conditions, Technologies, and Species.” Nature Biotechnology 36 (5): 411–20.

12. Castilletti, Concetta, Licia Bordi, Eleonora Lalle, Gabriella Rozera, Fabrizio Poccia, Chiara Agrati, Isabella Abbate, and Maria R. Capobianchi. 2005. “Coordinate Induction of IFN-Alpha and -Gamma by SARS-CoV Also in the Absence of Virus Replication.” Virology 341 (1): 163–69.

13. Chen, Nanshan, Min Zhou, Xuan Dong, Jieming Qu, Fengyun Gong, Yang Han, Yang Qiu, et al. 2020. “Epidemiological and Clinical Characteristics of 99 Cases of 2019 Novel Coronavirus Pneumonia in Wuhan, China: A Descriptive Study.” The Lancet 395 (10223): 507–13.

14. Cheung, Chung Y., Leo L. M. Poon, Iris H. Y. Ng, Winsie Luk, Sin-Fun Sia, Mavis H. S. Wu, Kwok-Hung Chan, et al. 2005. “Cytokine Responses in Severe Acute Respiratory Syndrome Coronavirus-Infected Macrophages in Vitro: Possible Relevance to Pathogenesis.” Journal of Virology 79 (12): 7819–26.

15. Chu, Hin, Jie Zhou, Bosco Ho-Yin Wong, Cun Li, Jasper Fuk-Woo Chan, Zhong-Shan Cheng, Dong Yang, et al. 2016. “Middle East Respiratory Syndrome Coronavirus Efficiently Infects Human Primary T Lymphocytes and Activates the Extrinsic and Intrinsic Apoptosis Pathways.” The Journal of Infectious Diseases 213 (6): 904–14.

16. Chu, Hin, Jie Zhou, Bosco Ho-Yin Wong, Cun Li, Zhong-Shan Cheng, Xiang Lin, Vincent Kwok-Man Poon, et al. 2014. “Productive Replication of Middle East Respiratory Syndrome Coronavirus in Monocyte-Derived Dendritic Cells Modulates Innate Immune Response.” Virology 454-455 (April): 197–205.

17. Delorey, Toni M., Carly G. K. Ziegler, Graham Heimberg, Rachelly Normand, Yiming Yang, Åsa Segerstolpe, Domenic Abbondanza, et al. 2021. “COVID-19 Tissue Atlases Reveal SARS-CoV-2 Pathology and Cellular Targets.” Nature 595 (7865): 107–13.

18. Gómez-Rial, Jose, Maria José Currás-Tuala, Irene Rivero-Calle, Alberto Gómez-Carballa, Miriam Cebey-López, Carmen Rodríguez-Tenreiro, Ana Dacosta-Urbieta, et al. 2020. “Increased Serum Levels of sCD14 and sCD163 Indicate a Preponderant Role for Monocytes in COVID-19 Immunopathology.” Frontiers in Immunology 11 (September): 560381.

19. Gu, Jiang, Encong Gong, Bo Zhang, Jie Zheng, Zifen Gao, Yanfeng Zhong, Wanzhong Zou, et al. 2005. “Multiple Organ Infection and the Pathogenesis of SARS.” The Journal of Experimental Medicine 202 (3): 415–24.

20. Hoffmann, Markus, Hannah Kleine-Weber, Simon Schroeder, Nadine Krüger, Tanja Herrler, Sandra Erichsen, Tobias S. Schiergens, et al. 2020. “SARS-CoV-2 Cell Entry Depends on ACE2 and TMPRSS2 and Is Blocked by a Clinically Proven Protease Inhibitor.” Cell 181 (2): 271–80.e8.

21. Huang, Chaolin, Yeming Wang, Xingwang Li, Lili Ren, Jianping Zhao, Yi Hu, Li Zhang, et al. 2020. “Clinical Features of Patients Infected with 2019 Novel Coronavirus in Wuhan, China.” The Lancet 395 (10223): 497–506.

22. Huang, Cheng, Naoto Ito, Chien-Te K. Tseng, and Shinji Makino. 2006. “Severe Acute Respiratory Syndrome Coronavirus 7a Accessory Protein Is a Viral Structural Protein.” Journal of Virology 80 (15): 7287–94.

23. Huang, Cheng, C. J. Peters, and Shinji Makino. 2007. “Severe Acute Respiratory Syndrome Coronavirus Accessory Protein 6 Is a Virion-Associated Protein and Is Released from 6 Protein-Expressing Cells.” Journal of Virology 81 (10): 5423–26.

24. Ito, Naoto, Eric C. Mossel, Krishna Narayanan, Vsevolod L. Popov, Cheng Huang, Taisuke Inoue, Clarence J. Peters, and Shinji Makino. 2005. “Severe Acute Respiratory Syndrome Coronavirus 3a Protein Is a Viral Structural Protein.” Journal of Virology 79 (5): 3182–86.

25. Jones, Terry C., Guido Biele, Barbara Mühlemann, Talitha Veith, Julia Schneider, Jörn Beheim-Schwarzbach, Tobias Bleicker, et al. 2021. “Estimating Infectiousness throughout SARS-CoV-2 Infection Course.” Science 373 (6551). https://doi.org/10.1126/science.abi5273.

26. Kamitani, Wataru, Krishna Narayanan, Cheng Huang, Kumari Lokugamage, Tetsuro Ikegami, Naoto Ito, Hideyuki Kubo, and Shinji Makino. 2006. “Severe Acute Respiratory Syndrome Coronavirus nsp1 Protein Suppresses Host Gene Expression by Promoting Host mRNA Degradation.” Proceedings of the National Academy of Sciences of the United States of America 103 (34): 12885–90.

27. Kowal, Krzysztof, Richard Silver, Emila Sławińska, Marek Bielecki, Lech Chyczewski, and Otylia Kowal-Bielecka. 2011. “CD163 and Its Role in Inflammation.” Folia Histochemica et Cytobiologica / Polish Academy of Sciences, Polish Histochemical and Cytochemical Society 49 (3): 365–74.

28. Lee, Jeffrey Y., Peter Ac Wing, Dalia S. Gala, Marko Noerenberg, Aino I. Järvelin, Joshua Titlow, Xiaodong Zhuang, et al. 2022. “Absolute Quantitation of Individual SARS-CoV-2 RNA Molecules Provides a New Paradigm for Infection Dynamics and Variant Differences.” eLife 11 (January). https://doi.org/10.7554/eLife.74153.

29. Lei, Xiaobo, Xiaojing Dong, Ruiyi Ma, Wenjing Wang, Xia Xiao, Zhongqin Tian, Conghui Wang, et al. 2020. “Activation and Evasion of Type I Interferon Responses by SARS-CoV-2.” Nature Communications 11 (1): 3810.

30. Liao, Mingfeng, Yang Liu, Jing Yuan, Yanling Wen, Gang Xu, Juanjuan Zhao, Lin Cheng, et al. 2020. “Single-Cell Landscape of Bronchoalveolar Immune Cells in Patients with COVID-19.” Nature Medicine 26 (6): 842–44.

31. Li, Jin-Yan, Ce-Heng Liao, Qiong Wang, Yong-Jun Tan, Rui Luo, Ye Qiu, and Xing-Yi Ge. 2020. “The ORF6, ORF8 and Nucleocapsid Proteins of SARS-CoV-2 Inhibit Type I Interferon Signaling Pathway.” Virus Research 286 (September): 198074.

32. Nelde, Annika, Tatjana Bilich, Jonas S. Heitmann, Yacine Maringer, Helmut R. Salih, Malte Roerden, Maren Lübke, et al. 2021. “SARS-CoV-2-Derived Peptides Define Heterologous and COVID-19-Induced T Cell Recognition.” Nature Immunology 22 (1): 74–85.

33. Ng, Kevin W., Nikhil Faulkner, Georgina H. Cornish, Annachiara Rosa, Ruth Harvey, Saira Hussain, Rachel Ulferts, et al. 2020. “Preexisting and de Novo Humoral Immunity to SARS-CoV-2 in Humans.” Science 370 (6522): 1339–43.

34. Ng, Lisa F. P., Martin L. Hibberd, Eng-Eong Ooi, Kin-Fai Tang, Soek-Ying Neo, Jenny Tan, Karuturi R. Krishna Murthy, et al. 2004. “A Human in Vitro Model System for Investigating Genome-Wide Host Responses to SARS Coronavirus Infection.” BMC Infectious Diseases 4 (September): 34.

35. Nouailles, Geraldine, Emanuel Wyler, Peter Pennitz, Dylan Postmus, Daria Vladimirova, Julia Kazmierski, Fabian Pott, et al. 2021. “Temporal Omics Analysis in Syrian Hamsters Unravel Cellular Effector Responses to Moderate COVID-19.” Nature Communications 12 (1): 4869.

36. Prebensen, Christian, Peder L. Myhre, Christine Jonassen, Anbjørg Rangberg, Anita Blomfeldt, My Svensson, Torbjørn Omland, and Jan-Erik Berdal. 2021. “Severe Acute Respiratory Syndrome Coronavirus 2 RNA in Plasma Is Associated With Intensive Care Unit Admission and Mortality in Patients Hospitalized With Coronavirus Disease 2019.” Clinical Infectious Diseases: An Official Publication of the Infectious Diseases Society of America.

37. Qin, Chuan, Luoqi Zhou, Ziwei Hu, Shuoqi Zhang, Sheng Yang, Yu Tao, Cuihong Xie, et al. 2020. “Dysregulation of Immune Response in Patients With Coronavirus 2019 (COVID-19) in Wuhan, China.” Clinical Infectious Diseases: An Official Publication of the Infectious Diseases Society of America 71 (15): 762–68.

38. RECOVERY Collaborative Group, Peter Horby, Wei Shen Lim, Jonathan R. Emberson, Marion Mafham, Jennifer L. Bell, Louise Linsell, et al. 2021. “Dexamethasone in Hospitalized Patients with Covid-19.” The New England Journal of Medicine 384 (8): 693–704.

39. RECOVERY Collaborative Group, Peter W. Horby, Marion Mafham, Leon Peto, Mark Campbell, Guilherme Pessoa-Amorim, Enti Spata, et al. 2021. “Casirivimab and Imdevimab in Patients Admitted to Hospital with COVID-19 (RECOVERY): A Randomised, Controlled, Open-Label, Platform Trial.” bioRxiv. medRxiv. https://doi.org/10.1101/2021.06.15.21258542.

40. Rothe, Camilla, Mirjam Schunk, Peter Sothmann, Gisela Bretzel, Guenter Froeschl, Claudia Wallrauch, Thorbjörn Zimmer, et al. 2020. “Transmission of 2019-nCoV Infection from an Asymptomatic Contact in Germany.” The New England Journal of Medicine 382 (10): 970–71.

41. Schroeder, Simon, Fabian Pott, Daniela Niemeyer, Talitha Veith, Anja Richter, Doreen Muth, Christine Goffinet, Marcel A. Müller, and Christian Drosten. 2021. “Interferon Antagonism by SARS-CoV-2: A Functional Study Using Reverse Genetics.” The Lancet. Microbe 2 (5): e210–18.

42. Schulien, Isabel, Janine Kemming, Valerie Oberhardt, Katharina Wild, Lea M. Seidel, Saskia Killmer, Sagar, et al. 2021. “Characterization of Pre-Existing and Induced SARS-CoV-2-Specific CD8+ T Cells.” Nature Medicine 27 (1): 78–85.

43. Schulte-Schrepping, Jonas, Nico Reusch, Daniela Paclik, Kevin Baßler, Stephan Schlickeiser, Bowen Zhang, Benjamin Krämer, et al. 2020. “Severe COVID-19 Is Marked by a Dysregulated Myeloid Cell Compartment.” Cell 182 (6): 1419–40.e23.

44. Silvin, Aymeric, Nicolas Chapuis, Garett Dunsmore, Anne-Gaëlle Goubet, Agathe Dubuisson, Lisa Derosa, Carole Almire, et al. 2020. “Elevated Calprotectin and Abnormal Myeloid Cell Subsets Discriminate Severe from Mild COVID-19.” Cell 182 (6): 1401–18.e18.

45. Song, Xiang, Wei Hu, Haibo Yu, Laura Zhao, Yeqian Zhao, Xin Zhao, Hai-Hui Xue, and Yong Zhao. 2020. “Little to No Expression of Angiotensin-Converting Enzyme-2 on Most Human Peripheral Blood Immune Cells but Highly Expressed on Tissue Macrophages.” *Cytometry. Part A: The Journal of the International Society for Analytical Cytology*, December. https://doi.org/10.1002/cyto.a.24285.

46. Stewart, Sheila A., Derek M. Dykxhoorn, Deborah Palliser, Hana Mizuno, Evan Y. Yu, Dong Sung An, David M. Sabatini, et al. 2003. “Lentivirus-Delivered Stable Gene Silencing by RNAi in Primary Cells.” RNA 9 (4): 493–501.

47. Trombetta, Amelia C., Guilherme B. Farias, André M. C. Gomes, Ana Godinho-Santos, Pedro Rosmaninho, Carolina M. Conceição, Joel Laia, et al. 2021. “Severe COVID-19 Recovery Is Associated with Timely Acquisition of a Myeloid Cell Immune-Regulatory Phenotype.” Frontiers in Immunology 12 (June): 691725.

48. Wang, Dawei, Bo Hu, Chang Hu, Fangfang Zhu, Xing Liu, Jing Zhang, Binbin Wang, et al. 2020. “Clinical Characteristics of 138 Hospitalized Patients With 2019 Novel Coronavirus-Infected Pneumonia in Wuhan, China.” JAMA: The Journal of the American Medical Association 323 (11): 1061–69.

49. Wang, Yeming, Dingyu Zhang, Guanhua Du, Ronghui Du, Jianping Zhao, Yang Jin, Shouzhi Fu, et al. 2020. “Remdesivir in Adults with Severe COVID-19: A Randomised, Double-Blind, Placebo-Controlled, Multicentre Trial.” The Lancet 395 (10236): 1569–78.

50. Weinreich, David M., Sumathi Sivapalasingam, Thomas Norton, Shazia Ali, Haitao Gao, Rafia Bhore, Jing Xiao, et al. 2021. “REGEN-COV Antibody Cocktail Clinical Outcomes Study in Covid-19 Outpatients.” bioRxiv. medRxiv. https://doi.org/10.1101/2021.05.19.21257469.

51. Wendisch, Daniel, Oliver Dietrich, Tommaso Mari, Saskia von Stillfried, Ignacio L. Ibarra, Mirja Mittermaier, Christin Mache, et al. 2021. “SARS-CoV-2 Infection Triggers Profibrotic Macrophage Responses and Lung Fibrosis.” Cell, November. https://doi.org/10.1016/j.cell.2021.11.033.

52. Wölfel, Roman, Victor M. Corman, Wolfgang Guggemos, Michael Seilmaier, Sabine Zange, Marcel A. Müller, Daniela Niemeyer, et al. 2020. “Virological Assessment of Hospitalized Patients with COVID-2019.” Nature 581 (7809): 465–69.

53. Worm, Sjoerd H. E. van den, Klara Kristin Eriksson, Jessika C. Zevenhoven, Friedemann Weber, Roland Züst, Thomas Kuri, Ronald Dijkman, et al. 2012. “Reverse Genetics of SARS-Related Coronavirus Using Vaccinia Virus-Based Recombination.” PloS One 7 (3): e32857.

54. Xiong, Yong, Yuan Liu, Liu Cao, Dehe Wang, Ming Guo, Ao Jiang, Dong Guo, et al. 2020. “Transcriptomic Characteristics of Bronchoalveolar Lavage Fluid and Peripheral Blood Mononuclear Cells in COVID-19 Patients.” Emerging Microbes & Infections 9 (1): 761– 70.

55. Xu, Hao, Liang Zhong, Jiaxin Deng, Jiakuan Peng, Hongxia Dan, Xin Zeng, Taiwen Li, and Qianming Chen. 2020. “High Expression of ACE2 Receptor of 2019-nCoV on the Epithelial Cells of Oral Mucosa.” International Journal of Oral Science 12 (1): 8.

56. Zeng, F. Y., C. W. M. Chan, M. N. Chan, J. D. Chen, K. Y. C. Chow, C. C. Hon, K. H. Hui, et al. 2003. “The Complete Genome Sequence of Severe Acute Respiratory Syndrome Coronavirus Strain HKU-39849 (HK-39).” Experimental Biology and Medicine 228 (7): 866–73.

57. Zhang, Q., P. Bastard, Z. Liu, J. Le Pen, M. Moncada-Velez, J. Chen, M. Ogishi, et al. 2020. “Inborn Errors of Type I IFN Immunity in Patients with Life-Threatening COVID-19.” Science 370 (6515). https://doi.org/10.1126/science.abd4570.

58. Zhou, Ziliang, Chunliu Huang, Zhechong Zhou, Zhaoxia Huang, Lili Su, Sisi Kang, Xiaoxue Chen, et al. 2021. “Structural Insight Reveals SARS-CoV-2 ORF7a as an Immunomodulating Factor for Human CD14+ Monocytes.” iScience 24 (3): 102187.

59. Zou, Xin, Ke Chen, Jiawei Zou, Peiyi Han, Jie Hao, and Zeguang Han. 2020. “Single-Cell RNA-Seq Data Analysis on the Receptor ACE2 Expression Reveals the Potential Risk of Different Human Organs Vulnerable to 2019-nCoV Infection.” Frontiers of Medicine 14 (2): 185–92.

